# Scarring hair follicle destruction is driven by the collapse of EGFR-protected JAK-STAT1-sensitive stem cell immune privilege

**DOI:** 10.1101/2023.10.11.561653

**Authors:** Karoline Strobl, Jörg Klufa, Regina Jin, Lena Artner-Gent, Dana Krauß, Philipp Novoszel, Johanna Strobl, Georg Stary, Igor Vujic, Johannes Griss, Martin Holcmann, Matthias Farlik, Bernhard Homey, Maria Sibilia, Thomas Bauer

## Abstract

The hair follicle stem cell niche is an immune-privileged microenvironment, characterised by suppressed antigen presentation, thus shielding against permanent immune-mediated tissue damage. In this study, we demonstrate the protective role of hair follicle-specific epidermal growth factor receptor (EGFR) from scarring hair follicle degeneration. Mechanistically, disruption of EGFR signalling generates a cell intrinsic hypersensitivity within the JAK-STAT1 pathway, compromising the immune privilege in the context of CD8 T cell and NK cell-mediated inflammation. Genetic depletion of either JAK1/2 or STAT1 or topical therapeutic inhibition of JAK1/2 restores the immune privilege and activates stem cells to resume hair growth in mouse models of epidermal and hair follicle specific EGFR deletion. Skin biopsies from EGFR inhibitor-treated and from EGFR-independent cicatricial alopecia patients indicate active STAT1 signalling within the hair follicles. Notably, a case study of folliculitis decalvans, characterised by progressive hair loss, scaling and perifollicular erythema, demonstrates successful treatment with systemic JAK1/2 inhibition. Our findings offer mechanistic insights and present a therapeutic strategy for addressing scarring hair follicle destruction associated with EGFR-inhibitor therapy and cicatricial alopecia.

## Introduction

Hair serves an evolutionarily essential purpose in mammals by acting as a sensor, thermo-regulator and physical shield against external threats (*1*). Hair follicles, however, also represent a vulnerable portal within the epidermal barrier. Anatomical or immunological dysfunction of this unit can be exploited by microbes leading to hair follicle inflammation and the consecutive destruction of this essential body structure (*2*).

To counter tissue destruction during inflammatory insults, hair follicles have evolved an immune privilege (IP) mechanism (*3, 4*). This hair follicle stem cell IP is characterized by low expression of major histocompatibility complex class I (MHC I) for self-tolerance, up-regulation of “no danger” signals such as CD200, generation of an immunosuppressive microenvironment via TGF-β secretion, and recruitment of regulatory T cells, Trem2^+^ macrophages, and invariant natural killer (NK) T cells (*5–9*).

Autoimmunity mediated by cytotoxic T cells can lead to the destruction of the hair follicle bulb region, which consists of transiently amplifying cells during hair growth, resulting in reversible hair loss known as alopecia areata (AA) in humans (*10*). Several anti-inflammatory drugs, including corticosteroids, calcineurin inhibitors and janus kinase (JAK) inhibitors, have shown varying therapeutic effects in AA by targeting these T-cells (*11, 12*).

In contrast to AA, scarring (cicatricial) alopecia is characterized by the loss of the bulge region, which represents the hair follicle stem cell (HFSC) niche, leading to permanent and irreversible hair loss. Recent evidence implicates the collapse of the IP in HFSCs as a contributing factor in scarring alopecia types such as neutrophilic folliculitis decalvans and lymphocytic lichen planopilaris (*10, 13*). Hallmarks of these conditions include hyperproliferation and apoptosis of HFSCs, upregulation of MHC-I and MHC-II, reduced CD200 expression, and infiltration of various immune cells (*14, 15*). So far, the therapeutic options for scarring alopecia are limited by the lack of mechanistic understanding.

Patients with loss-of-function mutation of epidermal growth factor receptor (EGFR) or ADAM17, an EGFR ligands sheddase, as well as those undergoing long-term EGFR-inhibitor treatment during targeted cancer therapy, often experience severe chronic skin inflammation, which can be accompanied by scarring hair loss (*16–19*). The lack of effective treatments targeting the underlying causes of these cutaneous adverse events often leads to dose reduction or cessation of cancer therapy, compromising its efficacy (*20, 21*).

Recently, we have demonstrated the importance of EGFR-ERK signalling in maintaining barrier integrity and preventing bacterial invasion and dysbiosis during hair shaft eruption, revealing the initial structural and inflammatory trigger (*22–25*). We could show that the barrier defect can be rescued by re-establishing active ERK signalling independent of EGFR through transgenic over-expression of SOS under the K5 promoter (EGFR^Δep^ K5-SOS). In addition, the microbial arm of the skin inflammation can be reduced by broad spectrum antibiotic therapy (EGFR^Δep^ Abx). However, the molecular and immunological mechanisms driving and sustaining the chronic phase of the skin inflammation and hair loss remain enigmatic.

Here we show that EGFR protects against microbiota-driven destruction of the HFSC niche and subsequent scarring hair loss. Mechanistically, tissue destruction is initiated by the lack of EGFR-ERK signalling and driven by unleashed JAK-STAT1 activation in a cell autonomous manner. CD8^+^ T cells and NK cells expressing IFN-γ provoke disruption of the unguarded IP and its HFSC niche. Prophylactic and therapeutic JAK1/2 inhibition prevented the collapse of HFSC IP, restored hair growth, improved epidermal barrier function, and alleviated skin inflammation in epidermal and hair follicle specific EGFR deleted mouse models and a patient with previously untreatable folliculitis decalvans. Active STAT1 could be readily detected in patient samples during EGFR-inhibitor treatment and in different types of cicatricial alopecia.

These data represent mechanistic evidence of the HFSC intrinsic JAK-STAT1 cascade being responsible for exhausting the IP and implicates EGFR in securing its regulatory machinery. Our findings offer therapeutic strategies for effectively managing severe adverse events induced by EGFR inhibitors and scarring hair follicle destruction.

## Results

### Hair follicle-specific EGFR protects from microbiota-driven inflammatory hair follicle destruction

As previously published by our group, constitutive deletion of epidermal EGFR using K5-cre (EGFR^Δep^) results in epidermal barrier disruption and skin inflammation (*22*). Interestingly, the reduction of the inflammation by transgenic K5-SOS expression and antibiotic therapy prevented visible hair loss between 2 and 5 months of age in EGFR^Δep^ mice (Fig. 1A). Additional FACS analysis revealed that the inflammation is responsible for losing the CD34 positive hair follicle stem cells (HFSCs) and that this precedes macroscopic hair loss (Fig. 1A, B and Fig. S1A for overall FACS gating strategy). The rescue of HFSCs during antibiosis is paralleled by a reduction of αβT cells (Fig. S1B). Most EGFR^Δep^ mice die within the first couple of weeks after birth due to the inflammation, which hinders the effective study of the late chronic inflammatory stage including hair loss (*22*). Considering this, we generated a hair follicle-specific EGFR deletion mouse model using the Egr2-cre line (EGFR^ΔEgr2^, Fig. 1C-G) (*26*). Efficient deletion of EGFR in the hair follicles and retained interfollicular epidermal EGFR expression was confirmed using immune fluorescence (IF) staining of EGFR in EGFR^ΔEgr2^ skin sections (Fig. 1C). Most importantly, these mice develop early hair follicle-specific inflammation, as described by Langerhans cell (LC) specific MHC-II up-regulation around the hair follicles and later-occurring, microbiota-dependent hair loss (Fig. 1D and E). The restriction of the inflammation to the hair follicles leads to a dramatically improved survival of these mice over the pan-epidermal deletion model, which enabled us to study the events leading to the late-stage chronic skin inflammation and the subsequent hair loss in detail (Fig. S1C). Chronological analysis revealed that the HFSC niche is established during morphogenesis and persists up to the first month after birth (Fig. 1F). However, starting with the first anagen hair cycle phase after morphogenesis, the HFSCs gradually disappeared over time with no detectable CD34 positive HFSCs remaining after 5 months concomitant with an epidermal influx of immune cells and visible hair loss (Fig. 1E and F). Moreover, we also observed a time-delayed size reduction of the entire hair follicle (Fig. 1G and Fig. S1D). These data demonstrate that hair follicle-specific EGFR protects from microbiota-driven inflammatory hair follicle destruction and hair loss.

**Fig. 1.**
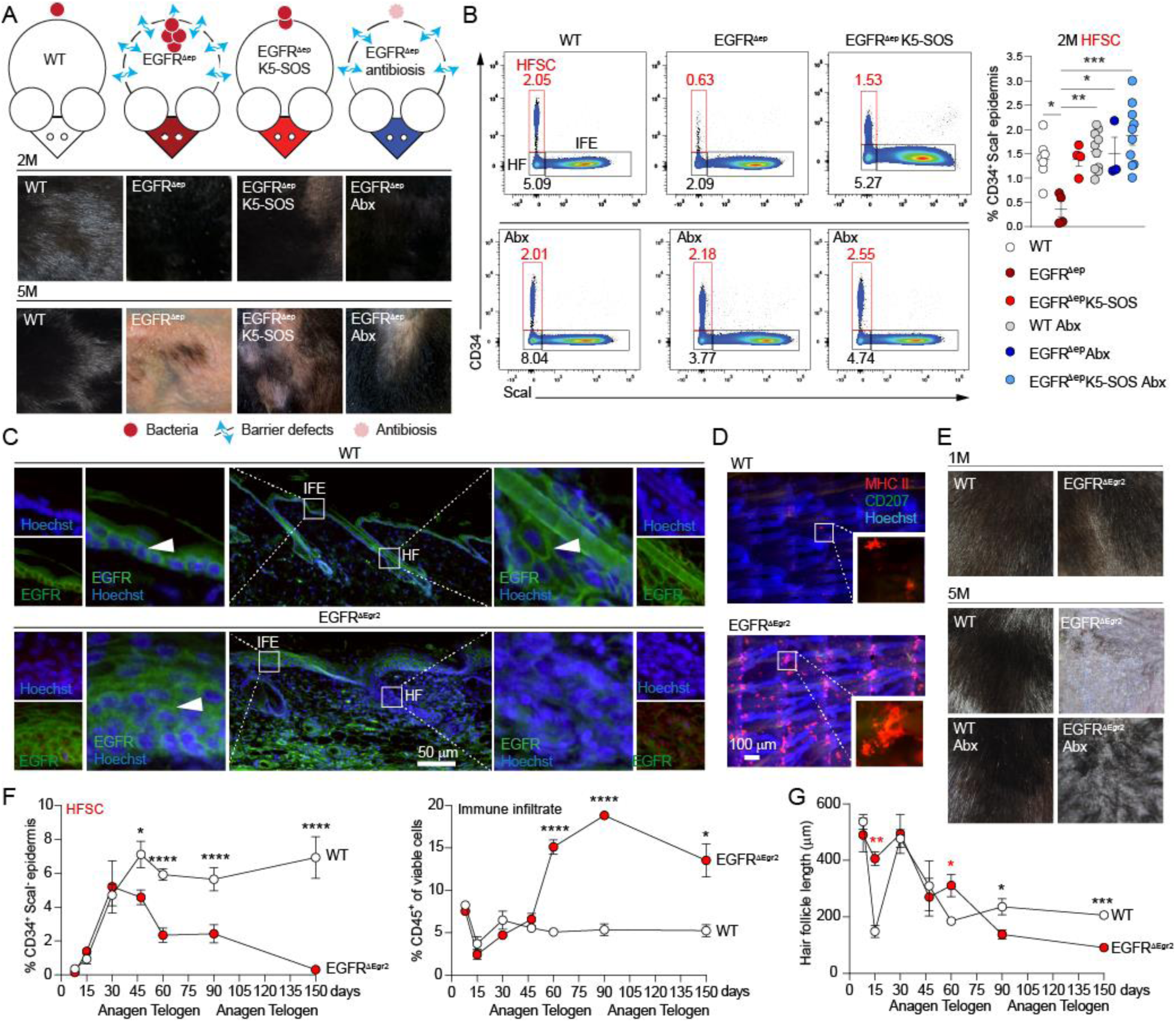
Hair follicle-specific EGFR protects from microbiota-driven inflammatory hair follicle destruction. **(A)** Graphical summary of mouse models used and representative pictures of their back-skin at 2 months (2M) and 5M of age. Wildtype (WT) mice have an intact barrier and normal microbiota. EGFR^Δep^ mice develop a skin barrier defect and microbial dysbiosis. EGFR^Δep^ K5-SOS mice have an intact barrier but develop microbial dysbiosis (*22*). Antibiotics (Abx, Cefazolin) treated EGFR^Δep^ mice are devoid of microbial inflammation while the initial barrier defect sustains. **(B)** Representative fluorescence-activated cell sorting (FACS) analysis of CD34^+^ Sca-I^-^ hair follicle stem cells (HFSC), hair follicles (HF) and interfollicular epidermis (IFE) of WT, EGFR^Δep^ and EGFR^Δep^ K5-SOS with or without antibiotic treatment at 2M of age and the % of HFSCs quantified. Detailed gating strategy and inflammatory status see also Fig. S1A and B. **(C)** Immunofluorescence (IF) staining of skin sections of EGFR (green) in EGFR^ΔEgr2^ mice and their respective WT controls. **(D)** LCs (CD207, green) and MHC-II^high^ expression (red) in epidermal sheets from tails of WT and EGFR^ΔEgr2^ mice. **(E)** Representative pictures of the hairy coat of EGFR^ΔEgr2^ at indicated time points and treatment. **(F)** Chronological change of the epidermal CD34^+^ HFSCs and CD45^+^ immune cells of EGFR^ΔEgr2^ and littermate WT controls as measured by FACS. **(G)** Chronological analysis of the hair follicle length as measured from hematoxylin and eosin (H&E) stained skin sections of WT or EGFR^ΔEgr2^ mice. Data acquisition see also in Fig. S1D. Data is presented in ±SEM, *P < 0.05, **P < 0.01, ***P < 0.001, ****P < 0.0001 by One-Way ANOVA with Tukey’s posthoc correction, n≥3.

### Hallmarks of scarring alopecia identified through RNA profiling of EGFR-deficient hair follicle bulge stem cells

Up to the first month of age, HFSCs can be detected in EGFR^ΔEgr2^ mice (Fig. 1F and Fig. 2A). In order to analyze their transcriptional status, we isolated these cells using the CD34 HFSC surface marker from WT and EGFR^ΔEgr2^ mice at 1 month and performed RNA sequencing (RNAseq). Efficient *Egfr* deletion was verified by RT-PCR from the sorted cells and *Egfr* is the most significantly down-regulated gene in the RNAseq dataset (Fig. S2A and B). Principal component analysis revealed the dramatic transcriptional differences between WT and EGFR^ΔEgr2^ HFSCs (Fig. S2C). Regulon analysis using DoRothEA revealed differential expression of transcription factors involved in cell cycle regulation (e.g. Foxm1, E2f2, E2f4 and Myc, Fig. 2B). Additionally, among the top 50 deregulated genes (DEG) in EGFR^ΔEgr2^ HFSCs was the downregulated *Bmp6*, implicated in HF quiescence and the upregulated proliferation-induced gene *Mki67* (Fig. S2D). These data prompted us to specifically look at proliferation, HFSC activation and quiescence genes, which revealed the hyper-proliferative status of the EGFR^ΔEgr2^ HFSCs compared to the WT situation (Fig. 2C). Next, we confirmed these findings using IF to detect the proliferating bulb region and Ki67 expressing bulge CD34 positive HFSCs (Fig. 2D). In line with this, shaved dorsal skin of 2-month-old EGFR^ΔEgr2^ mice revealed that EGFR^ΔEgr2^ mice remained in the anagen hair cycle phase (dark skin color) at 2 months of age as opposed to the light telogen skin of WT mice (Fig. S2E). Interestingly, gene set enrichment analysis (GSEA Wiki pathways), apart from proliferation, also identified signatures of apoptosis and fibrosis in these cells (Fig. 2E). Therefore, we performed IF analysis from 3-month-old EGFR^ΔEgr2^ mice, which confirmed apoptotic (Caspase 8^+^) HFSCs and Vimentin expression as a sign of fibrosis (Fig. 2F). After macroscopically visible hair loss at 5 months of age, skin sections of EGFR^ΔEgr2^ mice indicate follicular plugging and fibrotic hair structures, with a dramatic decline of hair follicles after 10 months of age (Fig. 2G and H and Fig. S2F). Taken together, we identified that during the chronic phase of skin inflammation, EGFR^ΔEgr2^ mice develop hallmarks of scarring hair follicle destruction.

**Fig. 2.**
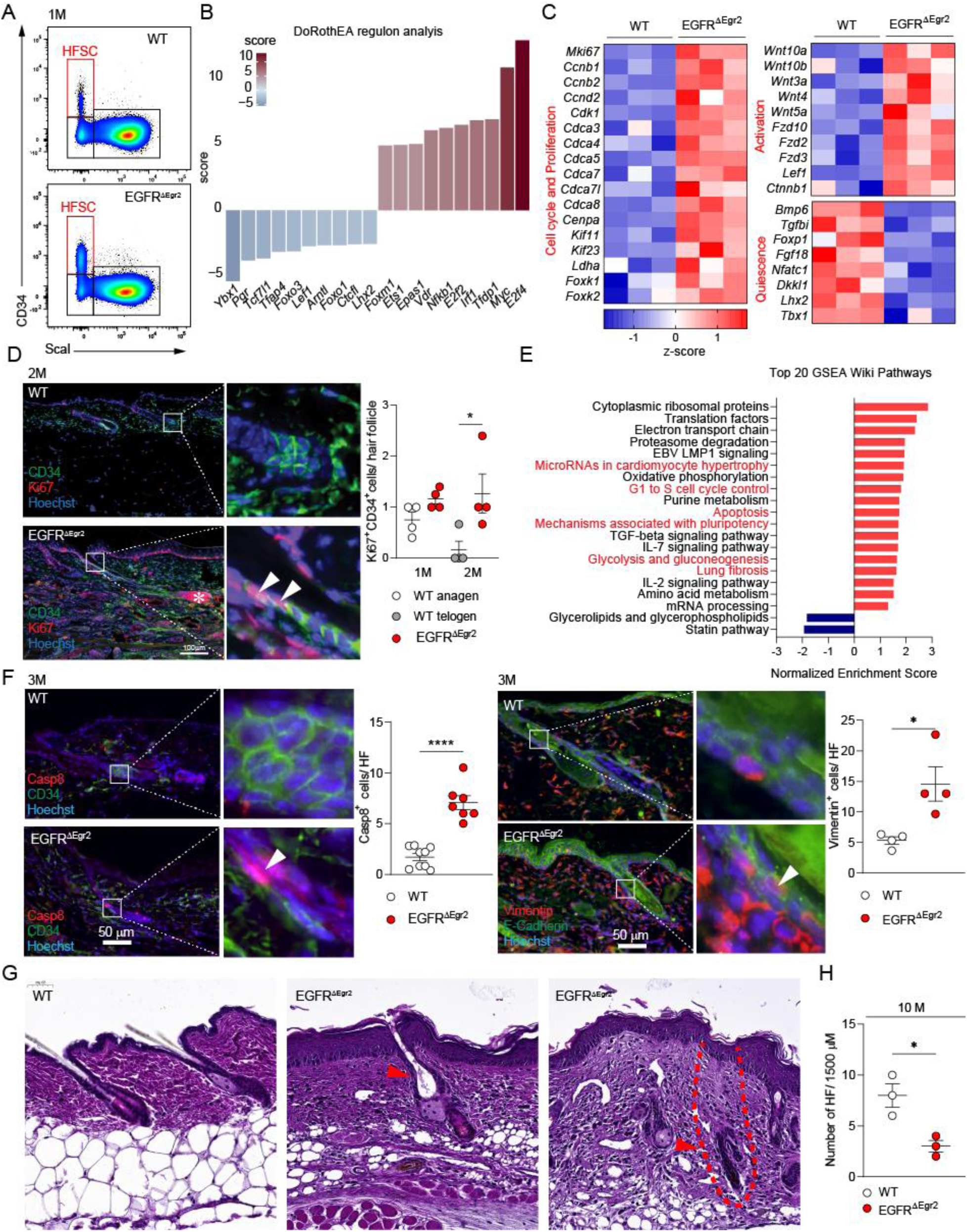
RNA profiling of EGFR-deficient hair follicle bulge stem cells identified hallmarks of scarring hair follicle destruction. **(A)** Representative FACS plot of sorting strategy for CD34^+^ScaI^-^ HFSCs (red box) of WT and EGFR^ΔEgr2^ mice at 1M of age. **(B)** DoRothEA transcription factor analysis of the HFSC RNAseq dataset. **(C)** Heatmap with z-scores of selected differentially expressed genes of the HFSC RNAseq dataset from WT and EGFR^ΔEgr2^ mice. **(D)** IF analysis and quantification of proliferation (Ki67, red) in HFSCs (CD34, green) from skin sections of WT and EGFR^ΔEgr2^ mice at 1M and 2M of age. **(E)** Top 20 gene set enrichment analysis (GSEA) WikiPathways from the HFSC RNAseq dataset. **(F)** Skin section IF staining of apoptosis (Caspase 8, red) and intermediate filament (Vimentin, red) expression in the hair follicle of WT and EGFR^ΔEgr2^ mice at 3M of age. **(G)** H&E histochemistry of WT and EGFR^ΔEgr2^ at 5M, red arrows indicate follicular plugging and a fibrotic hair follicle (indicated with a dotted line). **(H)** Number of hair follicle units per 1500 μm of H&E stained sections of WT and EGFR^ΔEgr2^ skin at 10M of age. Data is presented in ±SEM, *P < 0.05, ****P < 0.0001 by unpaired t-test, n≥3.

### Hair follicle intrinsic inflammatory JAK-STAT1 signaling induces immune privilege breakdown, hair follicle destruction and skin barrier disruption upon EGFR deletion

Next, we aimed to find the active signaling hubs in the HFSCs of EGFR^ΔEgr2^ mice. Cell signaling analysis from the HFSC RNAseq data set with PROGENy identified the activity of the inflammatory JAK-STAT and TNFα pathways (Fig. 3A). Previous studies indicated that the TNFα pathway does not play a prominent role in the inflammatory EGFR^Δep^ phenotype (*23, 27*). In line with the PROGENy pathway analysis from our EGFR^ΔEgr2^ HFSC RNA profiling, we detected a broad induction of the JAK-STAT signaling hub, including downstream effectors (e.g. interferon induced genes: *Ifi47*, *Ifi214* and *Ifitm3*) and the downregulation of its negative regulator suppressor of cytokine signaling 3 (*Socs3;* Fig. 3B, first panel). SOCS3 protein down-modulation could also be visualized in the hair follicles from skin sections from EGFR^ΔEgr2^ mice as compared to the WT (Fig. S3D). Notable, we also discovered a remarkable up-regulation of various JAK-STAT utilizing receptor complexes on EGFR^ΔEgr2^ HFSCs (Fig. 3B second panel). This prompted us to further investigate possible downstream effectors. MHC up-regulation has been linked to the JAK-STAT1 signaling cascade in keratinocytes (KCs) (*28, 29*). Indeed, we first recognized the upregulation of genes encoding for the MHC including its functional components (transporter associated with antigen processing: *Tap1* and *Tap2* genes) and regulators (Nod like receptor 5: *Nlrc5*) together with components of the immunoproteasome (e.g. proteasome subunit beta type-9, 8 and 10; Fig. 3B, third and fourth panel). MHC I and II protein up-regulation were confirmed on the cell membrane of epidermal cells from EGFR^ΔEgr2^ mice in the chronic inflammatory phase as compared to the WT situation using FACS (Fig. 3C and Fig. S3A for gating). Quiescent immune-privileged HFSCs have low MHC expression and are able to resist pro-inflammatory signals to prevent its up-regulation (*5*). Especially MHC-I antigen presentation on HFSCs is considered as breakdown of the immune privilege (*13*). EGFR^ΔEgr2^ HFSCs displayed MHC up-regulation on the RNA level and protein surface expression between 2 (postnatal day 15) and 4 (p30) weeks after birth (Fig. 3D and Fig. S3B). In line with immune privilege breakdown, we could additionally detect the downregulation of the anti-inflammatory surface proteins CD55 and CD200 in EGFR^ΔEgr2^ mice (Fig. S3C).

**Fig. 3.**
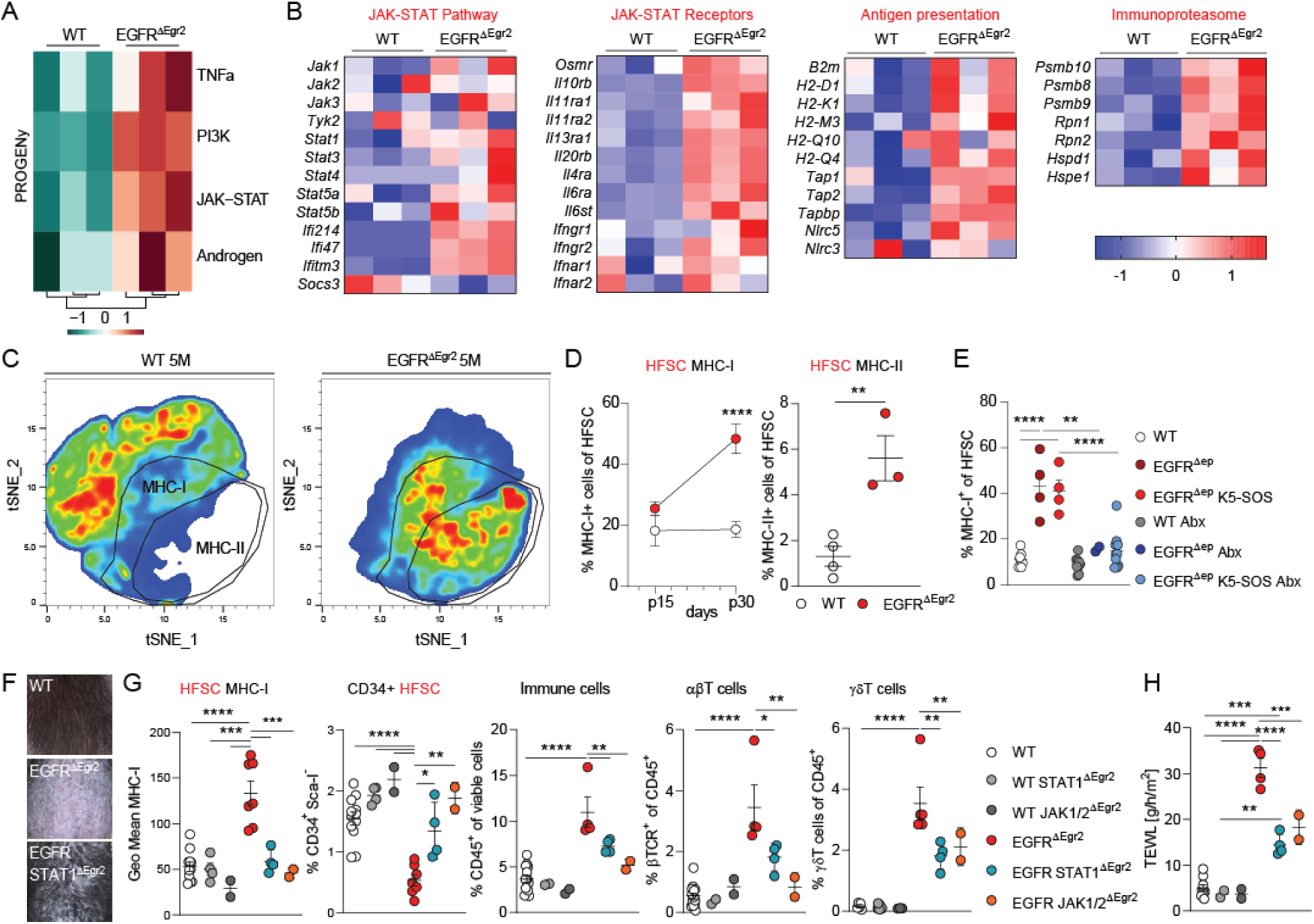
Hair follicle intrinsic JAK-STAT1 signaling induces immune privilege breakdown, hair follicle destruction and barrier disruption upon EGFR deletion. **(A)** PROGENy analysis identifying up-regulated signaling modules in the RNAseq dataset from sorted HFSCs from WT and EGFR^ΔEgr2^ mice. **(B)** Heatmap with z-scores of selected differentially expressed genes of the HFSC RNAseq dataset from WT and EGFR^ΔEgr2^ mice. **(C)** tSNE FACS plot of MHC-I and MHC-II expression among CD45^-^ epidermal cells of WT or EGFR^ΔEgr2^ at 5M. See also Fig. S3 for gating. **(D)** Percentage of MHC-I or MHC-II expression on CD34^+^ HFSCs by flow cytometry in WT and EGFR^ΔEgr2^ at 1M and **(E)** in WT, EGFR^Δep^, EGFR^Δep^ K5-SOS and antibiotics (Abx) treated mice at 2M. **(F)** Pictures of the hair phenotype of WT, EGFR^Δep^ and EGFR STAT1^ΔEgr2^ mice at the age of 5M. **(G)** The geometric mean of MHC-I, % of CD34^+^ HFSCs, total CD45^+^ cells, αβT cells and γδT cells among CD45^+^ cells by flow cytometry of EGFR STAT1^ΔEgr2^ and EGFR JAK1/2^ΔEgr2^ at 2M of age. **(H)** Transepidermal water loss (TEWL) measured on the back skin of WT, EGFR^ΔEgr2^, EGFR STAT1^ΔEgr2^ and EGFR JAK1/2^ΔEgr2^ mice. Data is presented in ±SEM, *P < 0.05, **P < 0.01, ***P < 0.001, ****P < 0.0001 by unpaired t-test or One-Way ANOVA with Tukey’s posthoc correction, n≥2.

Next, we utilized our EGFR^Δep^ K5-SOS/antibiosis model system to investigate the role of barrier immunity versus microbiota in influencing MHC regulation. HFSC-specific MHC-I surface expression was predominantly induced by the bacterial arm of the skin inflammation (Fig. 3E). In order to interfere with the hair follicle specific JAK-STAT1 cascade and investigate its involvement in the MHC expression status, skin function and inflammation, we crossed EGFR^ΔEgr2^ mice with either STAT1- or JAK1/2 floxed mice to generate double and triple hair follicle-specific knockout mice (Fig. 3F-H). Superficial hair loss was ameliorated in EGFR STAT1^ΔEgr2^ mice as compared to EGFR^ΔEgr2^ littermate controls (Fig. 3F). FACS analysis of the HFSCs revealed that their MHC-I expression is dependent on JAK-STAT1 signaling (Fig. 3G, HFSC MHC-I). Most importantly, HFSC destruction can be prevented by interrupting the cell-intrinsic JAK-STAT1 cascade (Fig. 3G, CD34+ HFSC). Additionally, we also observed an overall amelioration of the skin inflammation as measured by epidermal CD45^+^ immune cell infiltration and specifically αβ and γδT cell influx, which was paralleled by a reduction of the skin barrier defect (trans-epidermal water loss, TEWL) in the EGFR JAK1/2^ΔEgr2^ and EGFR STAT1^ΔEgr2^ mice, respectively (Fig. 3G and H). In summary, we conclude that the hair follicle-specific activation of the JAK-STAT1 cascade plays a dominant role in immune privilege collapse, hair follicle destruction and barrier immunity triggered by the lack of EGFR.

### Single-cell analysis revealed that immune privilege collapse and hair follicle destruction is driven by IFNγ-expressing NK and CD8 T cells

The initial trigger of the inflammation is a structural barrier breach at the hair follicle accompanied by microbial invasion, with the latter inducing JAK-STAT mediated MHC up-regulation, prolonging the barrier defect and driving αβ and γδT cell recruitment (Fig. 3E-G). This indicates a microbiota-dependent immune effector, which triggers and feeds into the hair follicle JAK-STAT1 cascade. Therefore, we next sought to investigate the immune infiltrate and the epidermal cytokine/chemokine milieu. Chronically inflamed, bald EGFR^ΔEgr2^ mice at 5 months of age show a dramatic shift in their epidermal immune cell compartment as compared to the homeostatic dendritic epidermal T cell (DETC) and LC networks in WT mice as measured by FACS (Fig. 4A and B). They lose these steady-state compartments and acquire an immune infiltrate dominated by αβ-, γδT cells and neutrophils (Fig. 4A and B).

**Fig. 4.**
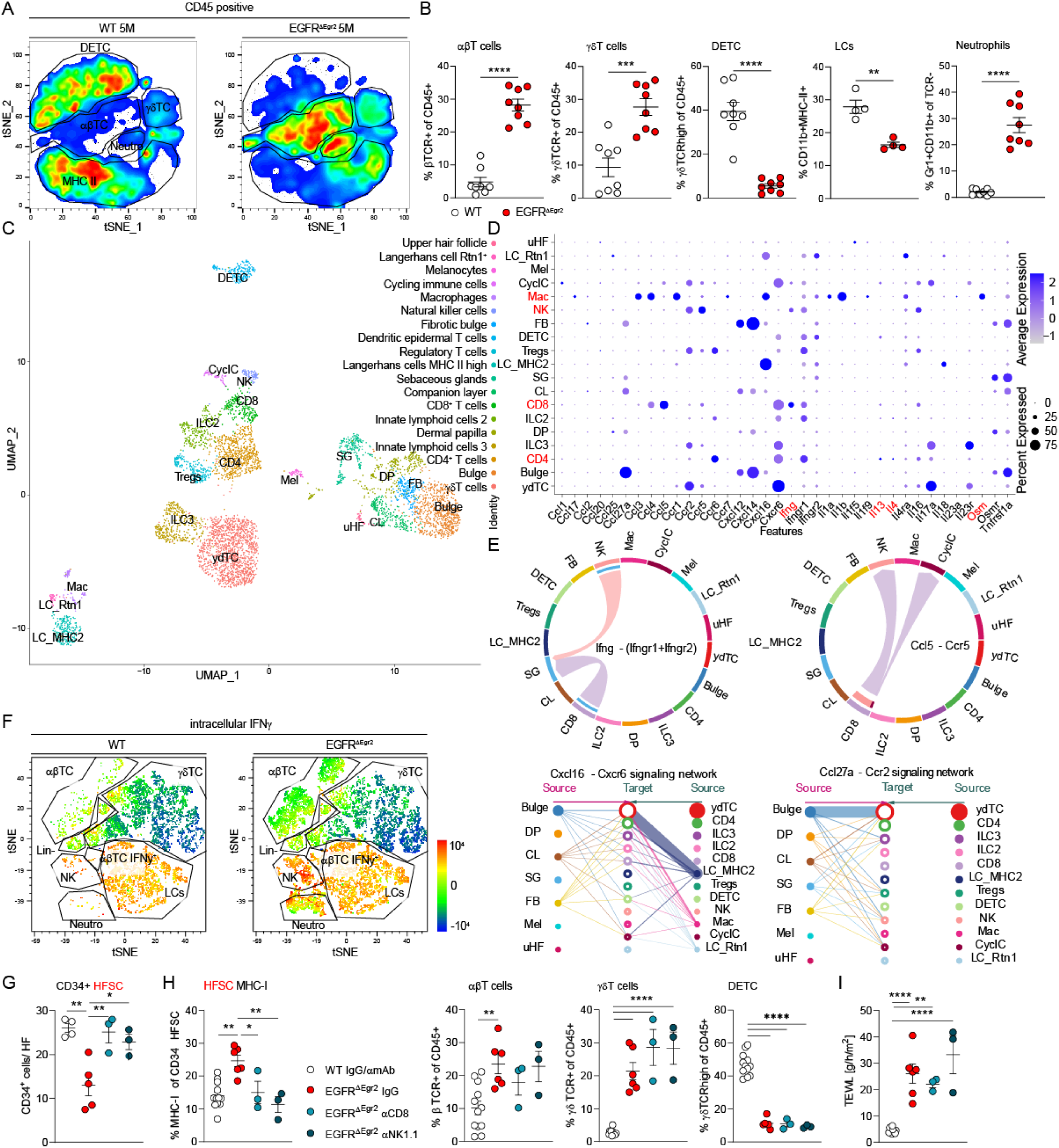
Single-cell analysis revealed that immune privilege collapse and hair follicle destruction is driven by IFNγ expressing NK and CD8 T cells. **(A)** tSNE FACS plot of epidermal CD45^+^ immune cells of WT or EGFR^ΔEgr2^ at 5M. See Fig. S1A for gating strategy. **(B)** Quantification of indicated epidermal immune cells by flow cytometry. **(C)** UMAP of single-cell RNA sequencing analysis of epidermal CD45^+^ immune cells and CD45^-^ Sca-I^-^ hair follicle cells of EGFR^ΔEgr2^ at 3M of age, see also Supplementary Figure 4 for population definition strategy. **(D)** Relative expression levels of selected cytokines and chemokines as bubble plot in cell subsets (possible JAK-STAT inducers highlighted in red). **(E)** CellChat cell interaction analysis of the single-cell dataset. **(F)** tSNE FACS plot of the intracellular expression level of IFNγ expressing (positive in red) immune cells of WT or EGFR^ΔEgr2^ at 5M of age. **(G)** Quantification of CD34^+^ HFSCs of IF staining. **(H)** Percentage of MHC-I expression on HFSCs and % of αβT cells, γδT cells and dendritic epidermal T cells (DETC) among CD45^+^ cells by flow cytometry from WT and EGFR^ΔEgr2^ depleted of CD8 T cells or NK1.1 cells and their respective controls. **(I)** Transepidermal water loss (TEWL) in CD8 T cell or NK1.1 cell depleted WT and EGFR^ΔEgr2^ mice. Data is presented in ±SEM, *P < 0.05, **P < 0.01, ***P < 0.001, ****P < 0.0001 by unpaired t-test or One-Way ANOVA with Tukey’s posthoc correction, n≥3.

In order to expand the depth of this analysis, we performed single-cell RNA sequencing of the hair follicle and immune cell compartment of EGFR^ΔEgr2^ mice at 3 months of age, which confirmed the FACS data and additionally identified two types of innate lymphoid cells (ILCs), natural killer cells (NK cells) and allowed stratification of T-cells into CD4, CD8 and T regulatory cells (Tregs, Fig. 4C and Fig. S4A). Confirming our data from Fig. 2, we also observed the fibrotic transition of the bulge compartment (fibrotic bulge, FB) indicating scarring hair follicle destruction (Fig. 4C and Fig. S4A). We next used this dataset to map the chemokine and cytokine profile of the immune- and hair follicle cell compartments (Fig. 4D). This captured the multifaceted inflammatory microenvironment in EGFR^ΔEgr2^ mice, with prominent *Il17*-producing γδT cells and ILC3s, *Il4* and *Il13* producing CD4 Th2 cells and ILC2s, *Ifnγ* producing CD8 T-cells and NK cells and IL1α/β and OSM expressing macrophages. Given the importance of the JAK-STAT cascade in the hair follicles of EGFR^ΔEgr2^ mice, we next sorted out the possible JAK-STAT inducers IL4, IL13, IFN-γ and OSM among the cytokine profile and screened for their potential in epidermal MHC regulation using an *ex vivo* skin explant culture system (Fig. S4B). We observed a significant up-regulation of MHC I and II only with IFN-γ, whether or not the tissue was incubated with the EGFR-inhibitor erlotinib (Fig. S4B). Putative cell-cell interaction analysis using CellChat indicated that the only cellular sources of *Ifnγ* were the CD8^+^ T and NK cells (Fig. 4E). CellChat also revealed a complex chemokine signaling network capable of recruiting the various T cell subsets and NK cells with *Ccl5* interlinking CD8^+^ T cells and NK cells (Fig. 4E)(*30*). Interestingly, the gene signatures of the CD8 T cell and NK compartments cluster relatively close together (Fig. 4C and Fig. S4C). Intracellular FACS analysis for IFN-γ confirmed its up-regulation in infiltrating αβT cells and NK cells at the protein level (Fig. 4F).

In order to investigate the functional role of these cells during hair follicle destruction we used depletion antibodies for CD8 or NK1.1 *in vivo*. After confirming the cell deletion efficiency by the antibodies, we could show by FACS analysis, that both CD8 T cells and NK cells are influencing the HFSC immune privilege status and survivability (Fig. S4C, D and Fig. 4G). Interestingly, however, CD8 T cell and NK cell depletion did not impact the γδT cell compartment and did not ameliorate the epidermal barrier defect as observed with JAK1/2 and STAT1 deficient hair follicles (compare Fig. 3H and I with Fig. 4H and I). We, therefore, conclude that among the heterogeneous immune environment induced by prolonged EGFR deficiency, IFNγ-producing CD8 T cells and NK cells drive the hair follicle JAK-STAT dependent immune privilege decline and initiate the HFSC destruction.

### Therapeutic JAK inhibition re-initiates hair growth, rescues the immune privilege and ameliorates overall skin function and inflammation in EGFR^ΔEgr2^ and EGFR^Δep^ mice

We next tested the direct effect of IFN-γ on *in vitro* KCs, either from EGFR^Δep^ mice with their respective WT controls or WT mice treated with or without the EGFR-inhibitor erlotinib. MHC II induction was dramatically enhanced in EGFR-depleted or inhibited KCs as compared to WT induction and this expression could be inhibited by the clinically approved JAK1/2 inhibitor ruxolitinib (Fig. 5A and Fig. S5A). Encouraged by these results, we next applied the JAK1/2 inhibitor topically on 5-month-old bald EGFR^ΔEgr2^ mice (Fig. 5B). Although, these mice had only a rudimentary HFSC niche and already degraded hair follicles (see Fig. 2), we could, during the course of 28 days treatment, observe the induction of novel hair regrowth (Fig. 5B). This visible therapeutic effect was accompanied by the reduction of MHC surface expression and the re-appearance of the HFSCs (Fig. 5C and D). Interestingly, however, the re-activated HFSCs still had reduced expression levels of the mature stem cell surface marker CD34 but could be readily detected by the stem cell marker SOX9 (Fig. 5D and Fig. S5B). Prophylactic ruxolitinib treatment starting from 1 month old EGFR^ΔEgr2^ mice, however, prevented the loss of CD34 surface expression, similar to the JAK1/2 or STAT1 deletion in this model (Fig. S5C). In the therapeutic setting, the re-appearance of SOX9 positive HFSCs was concomitant with an increase in hair follicle length (Fig. 5E). Apart from novel hair re-growth, therapeutic JAK1/2 inhibition was able to restore the epidermal barrier function, normalize epidermal thickness and stopped αβT cell infiltration (Fig. 5F-H). Interestingly, JAK1/2 inhibition did not influence the γδT cell compartment, the neutrophil recruitment and the NK cells (Fig. S5D).

**Fig. 5.**
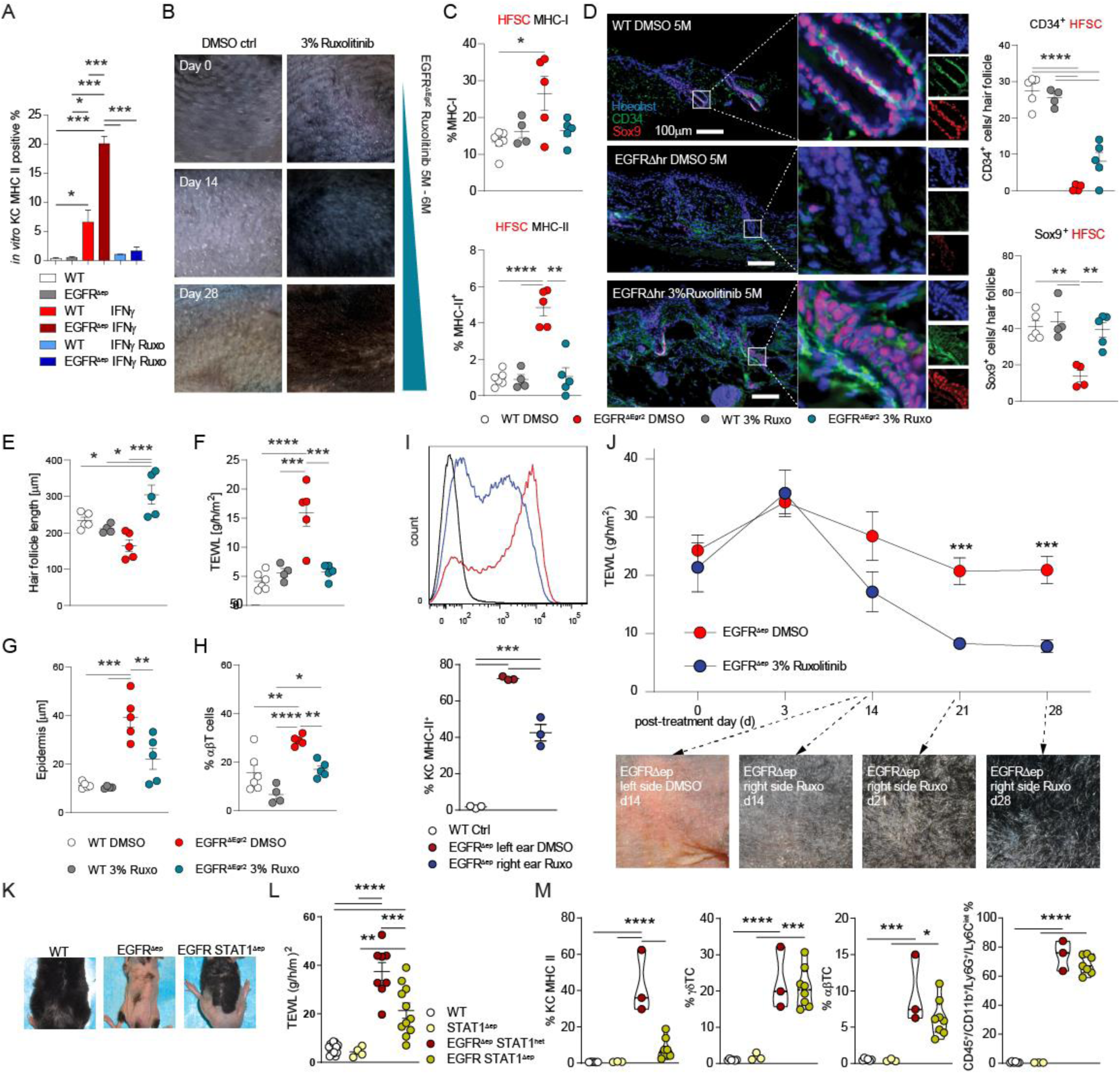
Therapeutic JAK inhibition re-initiates hair growth, re-installs the immune privilege and ameliorates overall skin function and inflammation in EGFR^ΔEgr2^ and EGFR^Δep^ mice. **(A)** MHC-II expression as analysed by flow cytometry of *in vitro* EGFR^Δep^ or WT primary murine keratinocytes treated with and without IFNγ and JAK1/2 inhibitor (ruxolitinib) as indicated. **(B)** Representative pictures of hair growth initiation in EGFR^ΔEgr2^ mice treated topically with DMSO (vehicle) or 3% ruxolitinib in DMSO daily at the age of 5M for 4 weeks. **(C)** Percentage of MHC-I or MHC-II expression in HFSCs by flow cytometry, **(D)** Immunofluorescence staining and quantification of CD34^+^ and Sox9^+^ HFSCs, **(E)** Hair follicle length as measured from H&E stained skin sections, **(F)** TEWL measurement, **(G)** Epidermal thickness measured from H&E stained skin sections and **(H)** % of αβT cells among CD45^+^ immune cells by flow cytometry from EGFR^ΔEgr2^ mice treated topically with DMSO (vehicle) or 3% ruxolitinib in DMSO daily at the age of 5M for 4 weeks and the respective WT controls **(I)** Percentage of MHC-II expression in keratinocytes by flow cytometry of EGFR^Δep^ or WT treated with DMSO or 3% ruxolitinib on the right and the left ear, respectively **(J)** TEWL and representative pictures of EGFR^Δep^ or WT over the 4-week treatment period with 3% ruxolitinib. **(k-m)** WT, EGFR^Δep^ and EGFR STAT1^Δep^ mice were analysed for their hairy coat **(K)**, TEWL **(L)** and for their inflammatory status by FACS for the indicated parameters, **(M).** Data is presented in ±SEM, **P < 0.01, ***P < 0.00, ****P < 0.0001 by One-Way ANOVA with Tukey’s posthoc correction, n≥3.

In order to extrapolate the mechanism and the therapeutic protocol to the full-scale skin inflammation model, we applied the JAK1/2 inhibitor to the right side (including the ear) of mice lacking EGFR in the complete epidermal compartment (EGFR^Δep^ mice) with the left side treated with vehicle control (Fig. 5I-J). Similar to the results from the EGFR^ΔEgr2^ hair follicle model, JAK1/2 inhibition reduced MHC expression, re-established skin barrier function and restored hair growth on the right side of the mice (Fig. 5I-J). Especially the restored epidermal barrier function and the hair regrowth indicates the successful preclinical therapeutic approach for treating the full blown skin inflammation. We next crossed EGFR^Δep^ mice with STAT1 floxed animals and could observe similar prophylactic effects as with the EGFR^ΔEgr2^ hair follicle model (compare Fig. 3H-J to Fig. 5K-M).

We could previously demonstrate that EGFR-controlled ERK signaling prevents the initial barrier disruption during novel hair shaft eruption or outgrowth (*22*). In order to investigate the involvement of the JAK cascade during this initial structural insult, we treated EGFR^Δep^ mice during hair eruption (starting from p6 until p19) topically with the JAK1/2 inhibitor (Fig. S 5E and F). As expected, this early prophylactic JAK1/2 inhibition did not impact the initial ERK dependent barrier breach, but only ameliorated MHC expression and prevented αβT cell influx. This indicates that the early barrier disruption is independent of JAK1/2 signaling and that hyper-activated JAK-STAT1 drives only the chronic phase of hair and skin inflammation. Taken together, we could demonstrate the effectiveness of topical JAK inhibition in reducing skin inflammation, restoring epidermal barrier function and reversing hair follicle destruction caused by prolonged EGFR dysfunction. These data represent pre-clinical evidence for the therapeutic potential of JAK inhibitors to manage adverse events during EGFR-inhibitor-targeted cancer therapy and reverse developing scarring hair follicle destruction.

### Phosphorylation of STAT1 in EGFR-inhibitor-treated patients and its relevance in cicatricial alopecia, along with the efficacy of JAK-inhibitor therapy in folliculitis decalvans

In order to confirm the most important hallmarks of our findings in patients, we next analyzed biopsies of EGFR-inhibitor treated squamous cell carcinoma (SCC) cancer patients. Comparison of pre- and post-EGFR-inhibitor treatment skin biopsies confirmed the breech of IP by epidermal MHC I upregulation (Fig. 6A). Furthermore, we detected the presence of phosphorylated STAT1 protein in clinical samples after EGFR inhibition, indicating its activation during the course of treatment (Fig. 6B). In line with our mouse model, phosphorylated STAT1 was elevated in the hair follicle and epidermis of patients with folliculitis decalvans, frontal fibrosing alopecia and lichen planopilaris (Fig. 6C and D and Fig. S6A).

**Fig. 6.**
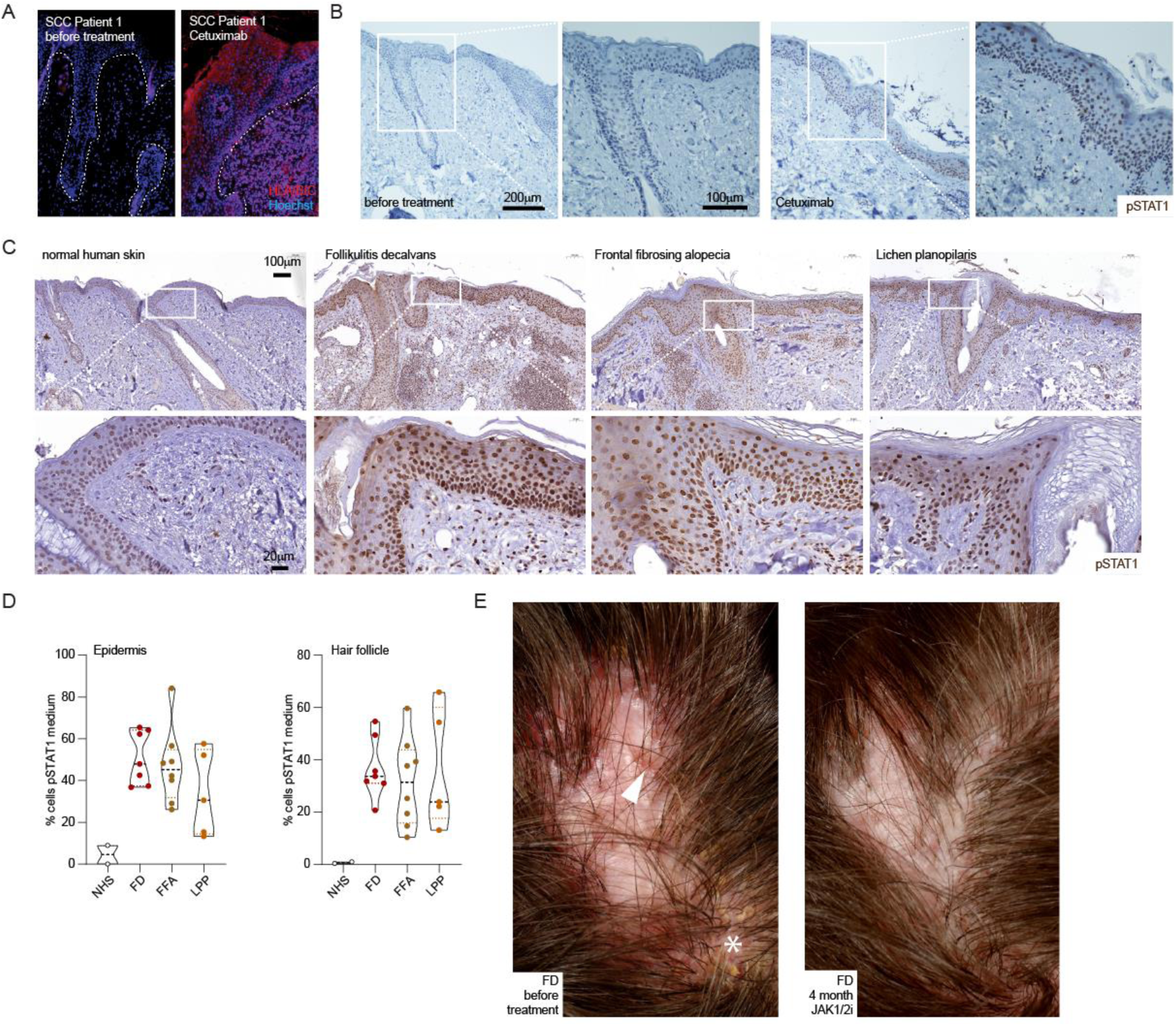
STAT1 activation in EGFR-inhibitor treated cancer patients and patients with scarring alopecia drives inflammation. **(A)** Human skin sections from biopsies of patients before EGFR-inhibitor treatment (left side) and after (right side) stained for MHC-I (red) and Hoechst (blue). Images are representative pictures from 2 independent patients. **(B)** Human skin sections stained immunohistochemically for phosphorylated STAT1 protein (pSTAT1) before EGFR-inhibitor treatment and after. Images are representative pictures from 2 independent patients. **(C)** Human skin sections from biopsies of patients before EGFR-inhibitor treatment (normal human skin) and patients with folliculitis decalvans (FD), frontal fibrosing alopecia (FFA) and lichen planopilaris (LPP, as indicated on top) stained for pSTAT1 immunohistochemically. Scale bar is 100μm for top panel and 20 μm for lower panel. **(D)** Quantification of pSTAT1 staining intensity as analysed from (C). Each dot represents an individual patient sample. See also in Fig. S6A. **(E)** Picture of clinical response after 4 month JAK-inhibitor therapy (oral Baricitinib) of folliculitis decalvans (arrowhead indicates representative perifollicular erythema, asterisk indicates area with yellowish, adherent scales before treatment: left picture).

Most importantly, we administered oral baricitinib, a JAK1/2 inhibitor, to a 31-year-old male patient diagnosed with folliculitis decalvans. Notably, no prior treatments yielded success for this patient over the course of three years. The histopathological examination revealed the presence of fibrotic hair follicles along with an inflammatory histiocytic and lymphocytic infiltrate (Fig. S6B). Noticeable improvements became apparent 4 weeks into JAK inhibitor therapy. Perifollicular erythema, yellowish and adherent scaling, and progressive scarring hair loss began to resolve and continued to improve over the subsequent 4 months (Fig. 6E). These positive clinical outcomes underscore the practical application of our research findings to human skin pathology. Moreover, they emphasize the significance of targeting the JAK-STAT1 signaling hub as a crucial therapeutic strategy in the context of EGFR inhibition and the treatment of human scarring alopecia.

## Discussion

The hairy coat is a survival necessity of mammals, as it protects from life-threatening outer influences like cold or ultraviolet radiation. In humans, reversible or irreversible hair loss, irrespective of the cause, severely affects the Quality-of-Life and psychological well-being of patients (*31*).

Using hair follicle-specific JAK1/2 and STAT1 knockout mice, we refined the spectrum of activity of the JAK inhibitors to HFSC-specific IP protection in scarring alopecia. As this condition is considered irreversible and current treatment regimens are only able to delay hair loss, it is of clinical importance that we were able to prophylactically prevent but also therapeutically reverse developing scarring hair follicle destruction using JAK inhibitors in mice and in a case study of human follicultis decalvans.

This suggests that not all HFSCs are destroyed simultaneously, but rather that the HFSC niche is constantly degraded and scarred over time. We observed a decline in HFSCs concomitant with the first anagen phase at 1 month of age and the start of visible hair loss from 3-month-old EGFR^ΔEgr2^ mice onward. Despite that, we were able to effectively regain hair growth up to 5 months of age. In older mice (e.g. 10 months of age), the complete scarring of the hair follicle then prevents therapeutic intervention indicating the irreversible loss of the stem cell niche. This represents evidence of a therapeutic window for treating scarring hair loss, although, in the various human conditions, it may differ greatly depending on the intensity of the inflammatory insult and the source and potency of the respective JAK-STAT1 activator.

It has been described that JAK inhibitors, although effective in some autoimmune diseases like rheumatoid arthritis or AA, do not necessarily have a clinically stable effect and after the cessation of the therapy, relapses are very common (*32, 33*). This might be reflective of the only temporary inhibiting effect on T cells. However, in scarring alopecia a more stable or longer lasting therapeutic effect could be possible in part due to the here-described multidirectional mode of action, which includes cell intrinsic restoration of the IP, blunting the recruiting and activation of the immune effectors, supporting anti-microbial immunity and by the restoration of skin barrier function. Additionally, other off-label case studies report successful JAK inhibitor therapy in cases of some scarring alopecia types (*34–36*). We now present the mechanistic explanation for these unbiased treatment attempts. Future clinical trials will disclose, whether JAK inhibitors can impact scarring alopecia in the long-term.

Upon EGFR inhibition, the initial immune response does not require the adaptive arm of the immune system. However, during the chronic phase, microbial dysbiosis leads to the recruitment of various T cell subsets to the epidermal compartment. Their cytokine profile indicates Th1, 17 and 2 responses, which are supported by their corresponding innate lymphoid cells, similar to what can be observed in atopic dermatitis (AD). Depleting CD8^+^ T cells expressing IFN-γ identified the driver of hair follicle-specific JAK-STAT1 activation and re-established the HFSC IP. Additionally, we identified NK cells as IFN-γ producers, which were also capable of disrupting the IP in our EGFR specific model system.

Interestingly, transcriptional profiling of the HFSCs revealed a broad up-regulation of JAK-STAT potent receptors. Among them the IL4- and IL13 receptors. Recent clinical case studies indicate that blocking IL4 and IL13 during AD-associated alopecia can reverse the hair loss indicating that various JAK-STAT inducers can feed into the HFSC intrinsic cascade (*37*). This additionally indicates that HFSC specific inhibition of the JAK-STAT1 cascade might act universally and is independent of the driving cytokines and its cellular source.

Therefore, it would be beneficial for patients with scarring alopecia, irrespective of the type, to screen for therapeutic markers highlighting JAK-STAT1 activity or JAK-inhibitor sensitivity to predict treatment efficiency or success. Common genetic signatures of cicatricial alopecia patients, identified fibrosis, immune cell pathways and susceptibility of frontal fibrosing alopecia with a locus containing the MHC-I region (*38, 39*). We could readily visualize the phosphorylation of STAT1 and the up-regulation of MHC-I on epidermal cells from patients under EGFR-inhibitor therapy and the pSTAT1 signature in patients with cicatricial alopeciás. These might represent possible candidates and readily available diagnostic markers for pre-screening patients with hair loss. However, more stable targets downstream of JAK-STAT1 like interferon induced transcription factors or soluble indicators measurable in liquid biopsies could help to effectively stratify patients in the future.

The molecular mechanism by which EGFR regulates the JAK-STAT1 sensitivity of epidermal cells might be ultimately driven by cell intrinsic receptor over-expression and simultaneous down-modulation of negative feedback mechanisms like SOCS3 as we could show here. EGFR, apart from regulating epidermal ERK and AKT signaling, is known to feed into the STAT3 cascade. Therefore, it is possible that the decline of epidermal EGFR mediated STAT3 signaling switches the balance in favor of STAT1 activation as it is described for STAT3 knockout conditions in various model systems including cancer cells (*40*).

It is intriguing to speculate that similar mechanisms of IP preservation in HFSCs, identified in this study, are also active in various EGFR-dependent solid tumors and that EGFR inhibition renders them more immunogenic, thereby supporting anti-cancer therapy efficiency. Indeed, it is a long-standing observation that the overall survival of cancer patients positively correlates to their EGFR-targeted anti-cancer therapy’s adverse events (*41*). Our study, however, indicates that the cell intrinsic JAK-STAT1 sensitivity introduced by the lack of EGFR also needs to be triggered and driven by the immune cell compartment, which opens room for improving or boosting the therapy effect by JAK activators.

Vice versa, further studies need to evaluate the feasibility of using JAK-inhibitors to treat EGFR-inhibitor induced adverse events, as JAK-inhibitors are predicted to rather support cancer growth by blunting the activity of cytotoxic T-cells. Therefore, topical application in this setting is crucial for the benefit of the patient and cancer therapy outcome.

Lastly, topical EGFR activation might represent a prophylactic strategy to strengthen the hair follicle IP and prevent disease onset in patients with genetic susceptibilities to scarring alopecia.

Taken together, our study identifies EGFR as cell intrinsic immune modulator, which tightens the strings on the JAK-STAT1 cascade to prevent inflammatory destruction of evolutionary important tissues like the hair follicle.

## Acknowledgments

We thank Sophie Huszarek and Lukas Ginzinger for excellent technical assistance. We thank Temenuschka Baykuscheva-Gentscheva for help with histology and Johannes Reisecker for his assistance in FACS sorting. The Core Facility Genomics at the Medical University of Vienna is acknowledged for carrying out the RNA sequencing analysis. We are grateful to Martina Hammer and the staff of the Department of Biomedical Research of the Medical University of Vienna for maintaining our mouse colonies.

## Author Contributions

K.S. designed the experiments, conducted experiments and wrote the paper; J.K., R.J., L-A.-G., P.N., M.H. conducted experiments; D.K. and R.J. analyzed data and performed statistical testing J.S., G.S., I.V., J.G., M.F. and B.H. designed the study, provided reagents, and revised the manuscript. M.S. supervised the project and provided funding. T.B. conceived and supervised the whole project, wrote the paper and provided the funding. All authors provided critical feedback to the manuscript.

## Data and materials availability

RNAseq datasets will be made available on Gene Expression Omnibus (GEO) database for publication. All other data are available in the main text or the supplementary materials.

## Methods

### Mice

EGFR^Δep^ and K5-SOS transgenic mice were generated as previously described in Lichtenberger et al.(*23*) and Klufa et al.(*22*). Egr2-Cre mice were purchased from The Jackon Laboratory. EGFR^ΔEgr2^ were generated by crossing Egr2-Cre with mice carrying conditional EGFR alleles EGFR^fl/fl^. STAT1 ^fl/fl^ were provided by Assoc. Prof. Priv.-Doz. Mag. Dr. Robert Eferl and generated by Wallner et al(*42*). JAK1^fl/fl^ and JAK2^fl/fl^ mice were provided by DI Dr Alexander Dohnal, Univ.-Prof. Dr. med.univ. Veronika Sexl and O.Univ.-Prof. Dr. med.vet. Matthias Müller and Univ.-Prof. Dr. Emilio Casanova. JAK2^fl/fl^ mice were generated as described in Wagner et al.(*43*) and Krempler et al.(*44*). EGFR STAT1^ΔEgr2^ or EGFR JAK1/2 ^ΔEgr2^ were generated by crossing EGR2-Cre and EGFR^fl/fl^, STAT1^fl/fl^, JAK1^fl/fl^ and JAK2^fl/fl^, respectively. Conditional knockout mice were kept with food and water ad libitum at the mouse facilities of the Medical University of Vienna and handled according to the standards and regulations approved by the animal experimental ethics committee and the Austrian Ministry of Science and Research (animal license number GZ BMWFW-66.009/0319-V/3b/2019). Corresponding to the Arrive guidelines (*45*), all experiments were designed to use the smallest number of animals, which are identified in the legends of the figures. Mice were allocated randomly to experiments and groups independent of sex. In all experimental set-ups, relevant treated and untreated wildtype and EGFR^ΔEgr2^ litter mates served as controls. Mice were treated with depletion recombinant antibodies i.p., topical pharmacological inhibitors and systemic antibiotics using the protocols below.

### TEWL measurement

TEWL of dorsal skin was measured with a Tewameter® TM 300 probe attached to the MDD4-display device (Courage + Khazaka) according to the manufacturer’s recommendations.

### Antibiotic treatment

The drinking water of mouse cages was supplemented with cefazolin (0.5g/L) (Astro Pharma) and replaced twice weekly. For increased cleanliness, cages were changed twice weekly.

### Depletion antibody treatment

For *in vivo* depletion, mice were administered i.p. twice weekly with 400 µg *InVivoMab* anti-NK1.1 (clone PK136), 300 µg *InVivoMab* anti-CD8α (clone 2.43, BioXCell) or *InVivoMab* rat IgG2b isotype control (clone LTF-2, BioXCell) in respective concentrations for 4 weeks.

### Topical Ruxolitinib treatment

Mice were treated topically daily with 100 µl 3% Ruxolitinib (LC Labs, R-6688) or vehicle control only (DMSO) for 4 weeks.

### Flow cytometric analysis of lymph node single-cell suspensions

Skin-draining lymph nodes were mechanically homogenized and filtered through a 70μm cell strainer to achieve a single-cell suspension in 2% FBS in PBS. The single cell suspension was blocked in FC-block CD16/32 (1:100, Biolegend) and subsequently incubated in Zombie Aqua viability stain (1:200, Biolegend for 15 min at room temperature. After another washing step, cells were resuspended in 50 μL staining buffer with the extracellular antibodies (see Table) and incubated for 30 min on ice. For the intracellular staining, cells were incubated with 50 μL of FOXP3 Fix/Perm (eBioscienceTM) working solution for 30 min at room temperature. Then the fixed solution was washed with 200 μL of FOXP3 permeabilisation buffer (eBioscienceTM), followed by the staining of FOXP3-PE in 50 μL of the same buffer for 60 min at room temperature. Cells were recorded using an LSR-II flow cytometer (BD Biosciences) and analyzed using FlowJo software.

### Fluorescence-activated cell sorting of epidermal single-cell suspensions

Harvested mouse ears were split into the dorsal and ventral sides and incubated in 0.8% trypsin (Gibco, Thermo Fisher Scientific) in PBS at 37℃ for 45 min. Dorsal skin was placed on 0.25% trypsin in DMEM and incubated at 4℃ overnight. The epidermis was separated from the dermis, further, digested in 250 µg/ml DNAseI (Sigma Aldrich) for 30 min at 37°C, and washed and filtered using a 70 µm cell strainer.

Single-cell suspensions were subsequently blocked with FC-block CD16/32 (Biolegend) and either stained with extracellular fluorescent antibodies at 4°C for 30 min and prior flow cytometric analysis SYTOX™ Blue Dead Cell Stain (1:1000, Thermo Fisher Scientific) was added. For intracellular staining of epidermal single-cell suspensions, cells were stained with Zombie Aqua viability stain (1:200, Biolegend) for 15 min at room temperature, following an extracellular antibody staining mix. Next, cells were fixed in 3% Formalin solution (Sigma Aldrich) in PBS for 10 minutes and permeabilized in Perm/Wash buffer (BD, 1:10) for 20 min at room temperature. Fluorescent antibodies for intracellular cytokines were added to Perm/Wash buffer and incubated for 30 minutes on ice. Cells were recorded using an LSR-II flow cytometer (BD Biosciences) or Cytek™ Aurora and analyzed using FlowJo software.

### Bulk RNA sequencing

Using the protocol above for Fluorescence-activated cell sorting of epidermal single cell suspensions, viable CD45^-^CD34^+^Sca-I^-^ cells were sorted from epidermal single cell suspensions of 1-month-old EGFR^ΔEgr2^ (n=3) and littermate controls (n=3) into Trizol LS Reagent (Thermo Fisher Scientific) using a FACS Aria III. RNA was extracted with the column-based miRNeasy Micro Kit (Qiagen) according to the manufacturer’s protocol. cDNA synthesis was performed with Superscript IV Reverse Transcriptase (Thermo Fisher Scientific). The main targets were confirmed by real-time PCR using SYBR-Green (Thermo Fisher Scientific). Samples were sent to Novogene, UK for quality control, library preparation and sequencing using Novaseq PE150. The kallisto pipeline was used to quantify the counts from the raw sequencing data (*46*). Differential gene expression analysis was performed using DESeq2 and Gene Set Enrichment Analysis (*47*).

### Single-cell RNA sequencing

Using the protocol above for Fluorescence-activated cell sorting of epidermal single cell suspensions, 12.000 viable CD45^-^Sca-I^-^ cells and 12.000 viable CD45^+^ were sorted from pooled epidermal single cell suspensions of 3-month-old EGFR^ΔEgr2^ (n=4) into a 0.04% BSA solution using a BD FACSMelody™ Cell Sorter (Biosciences). Single-cell cDNA libraries were generated using the droplet-based Chromium Next GEM Single Cell 5’ Kit v2 (Cat.nr. 1000265, 10x Genomics) following manufacturer’s instructions. Libraries were sequenced on insert machine name. The CellRanger pipeline version 7.0.0 was used to process raw sequencing files and produce feature-barcode matrices which were further analyzed in using several R packages. Most analyses were carried out using functions from the Seurat package(*48*) unless otherwise noted. First, ambient RNA contamination, which is commonly observed in droplet-based scRNA-seq methods, was corrected by using the SoupX package(*49*). Corrected count matrices were then imported into the standard Seurat workflow which includes initial quality control steps and filtering, cell cycle scoring, normalization, dimensionality reduction, clustering and visualization. Doublets were identified using the scDblFinder package(*50*). For the downstream analysis we included 5291 cells which passed quality control parameters (singlets, number of genes ≥ 300, number of UMIs ≥ 500, percentage of mitochondrial reads < 5%). Cell cycle scoring and the difference between G2M and S phase scores (“CC.Difference”) were calculated. Normalization was carried out using the SCTransform function with CC.Difference and percentage of mitochondrial reads set as variables to regress. For dimensionality reduction and visualization purposes, we used the RunPCA and RunUMAP functions of Seurat. For clustering and cluster marker identification, we used Seurat’s FindNeighbours, FindClusters and FindAllMarkers functions. Clusters were manually annotated using published single cell datasets of mouse skin(*51*). Predicted ligand-receptor interactions were determined with the CellChat package(*52*).

### *Ex-vivo* skin explant

Fresh mouse ears were split and floated dermal side down on 1ml RPMI medium containing 10% FBS and in a 24-well plate at 37°C for 48h. Medium was supplemented with 120 ng/mL INFγ, 20 ng/mL IL-4 (Preprotech), 20 ng/mL OSM or 20 ng/mL IL-13 (Bio-Techne). Analysis was performed using the protocol above for fluorescence-activated cell sorting of epidermal single-cell suspensions.

### Keratinocyte culture

Epidermal single-cell suspensions were isolated as previously described for fluorescence activated cell sorting and cultured on fibronectin-coated dishes in keratinocytes growth medium 2 (Promo2) supplemented with 0.02 mM CaCl2 (Promo Cell), growth supplements (1:40) and Penicillin/ Streptavidin (1:100, Sigma). 80% confluent cells were treated with Erlotinib (10 nM; Santa Cruz Biotechnology) for 24 hours. Next, the medium was supplemented with 120 ng/mL INFγ, (Peprotech) and 400 nM Ruxolitinib (LC Labs, R-6688).

### Full skin protein lysates and Luminex

Snap-frozen dorsal skin was added to RIPA lysis buffer, supplemented with Protease Inhibitor Cocktail (Roche) and homogenized using a Precellys 24 homogenizer (Bertin). Skin lysates were centrifuged at 14,000 x g for 15 min at 4°C to remove cell debris. For protein quantification, lysates were thawed on ice and subjected to Bradford protein quantification, according to the manufacturer’s protocol (Biorad). 70 – 100 μg total protein was used for each assay. Multiplex Luminex assays (Thermo Fisher Scientific) were performed according to the manufacturer’s recommendations and measured on a Luminex MAGPIX System using the xPONENT Software.

### Histological immunofluorescence and immunohistochemistry analysis

Dorsal skin was fixed in 4% PFA, embedded in a paraffin block and cut into 4 µm sections. After dewaxing and rehydration, sections were stained with hematoxylin and eosin (H&E), according to standard procedures of Papanicolaou. For immunofluorescence stainings, skin samples were heated in antigen retrieval solution using Dako Target Retrieval Solution (dilution 1:10, pH= 6 or pH=9). For immunohistochemistry, samples were treated with 3% H2O2 before blocking. All samples were blocked with 5% horse or goat serum in 2% BSA TBS-T for 1 hour in a humidified slide chamber at room temperature and stained with primary antibodies at 4°C overnight. The next day, slides were rinsed and incubated with an appropriate secondary fluorescence antibody (1:400) and Hoechst (Sigma Aldrich) or with Signal stain Boost IHC Detection reagent (HRP) for 2h in a dark humidified slide chamber. Tissue sections were mounted, and immunofluorescence pictures were taken using a Nikon Eclipse i80 microscope. For immunohistochemistry, skin tissues was stained with a DAB kit and hematoxylin. Slides were scanned and analysed using Definiense software.

### Human skin specimens and proof-of-concept treatment

Paraffin-embedded biopsy samples from folliculitis decalvans (n=7), frontal fibrosing alopecia (n=8) and lichen planopilaris (n=5) patients were obtained from the biobanks of the Departments of Dermatology at the University Hospital Düsseldorf and the Medical University of Vienna (Ethics approval ID 2016075402, Heinrich-Heine University, 40225 Duesseldorf, Germany; Ethics approval 1354/2021, Medical University of Vienna). Additionally, skin biopsies were available from two patients from the Department of Dermatology Rudolphstiftung Hospital (Vienna, Austria) who received cetuximab for the treatment of inoperable SCC and gave consent for the retrospective use of their data and samples. See more details in Klufa and Bauer et al (*22*). Moreover, we present a case with a proof-of-concept-treatment of a 31-year-old patient who experienced for 3 years inflammatory lesions on the scalp with progressive scaring alopecia as signs of a Folliculitis decalvans (Fig. 6e and Fig. S 6B; written consent of the patient was obtained for each photography). Multiple previous therapies, including anti-inflammatory and antibiotic measures, have been unsuccessful thus far. A shiny scar plate measuring approximately 5×3 cm was found in the parieto-occipital area of the capillitium. Multiple tufts of hair with perifollicular erythema and yellowish, adherent scaling were also present in the surrounding area. There were no relevant pre-existing conditions, and the family history was negative for other inflammatory skin diseases. Laboratory tests conducted before initiating an oral treatment with the JAK1/2 inhibitor baricitinib (4 mg/day) showed no abnormalities. A significant clinical improvement was observed already after 4 weeks of treatment and continued in the following months. The JAK inhibitor was well tolerated throughout the treatment period.

### Statistics

Statistical analyses were performed with GraphPad Prism 8.0 software. Data were tested for normal distribution by the Shapiro-Wilk test. In cases of normal distribution Student’s unpaired two-tailed t-test for comparisons of two groups or parametric One-way ANOVA analysis with Tukey’s pairwise comparisons were used to compare more than two groups. Experiments were repeated independently at least two times with similar results. Dot plots depict biological replicates unless otherwise stated. (*P < 0.05, **P < 0.01, ***P < 0.001, and ****P < 0.0001). The data are shown as mean ± SEM.

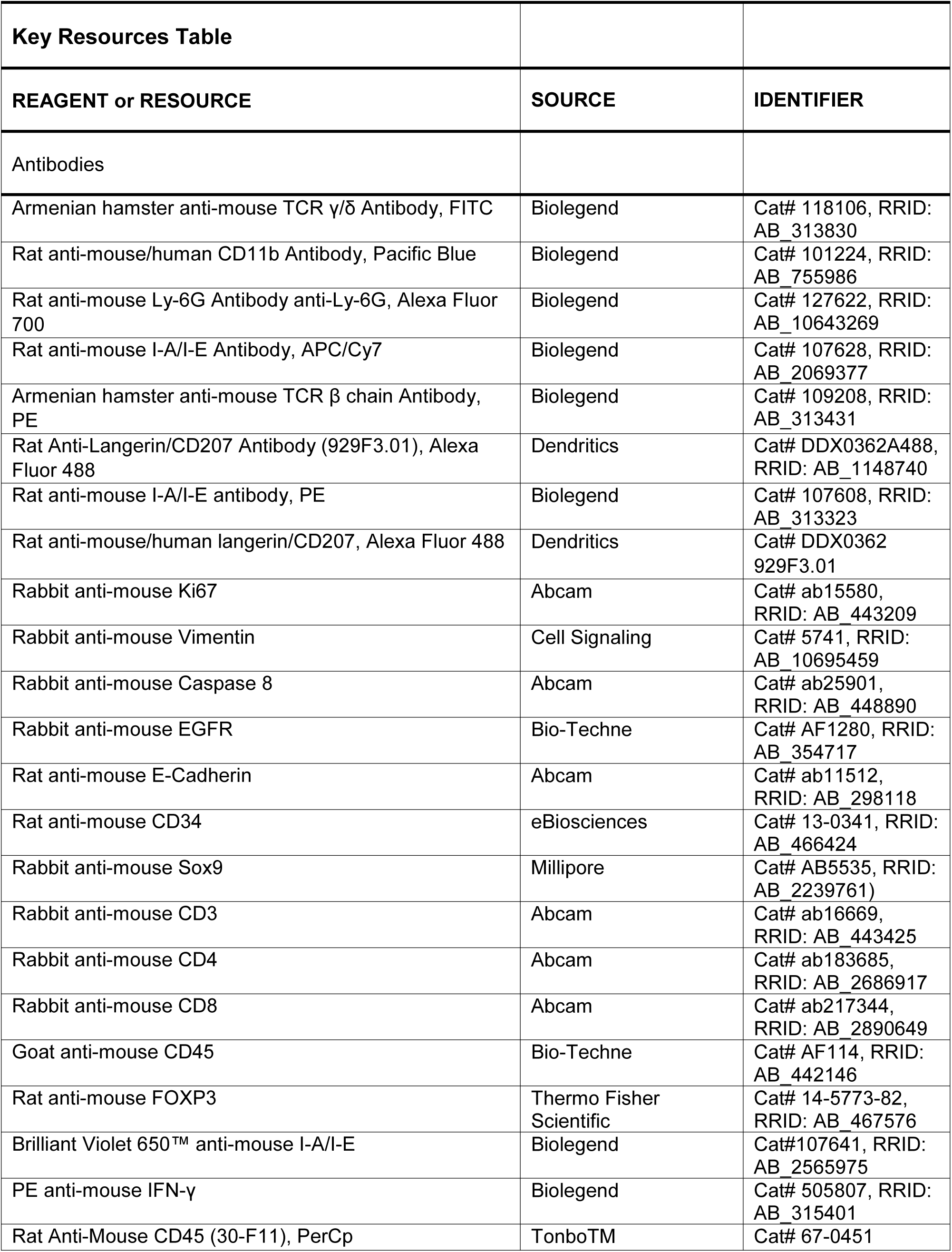
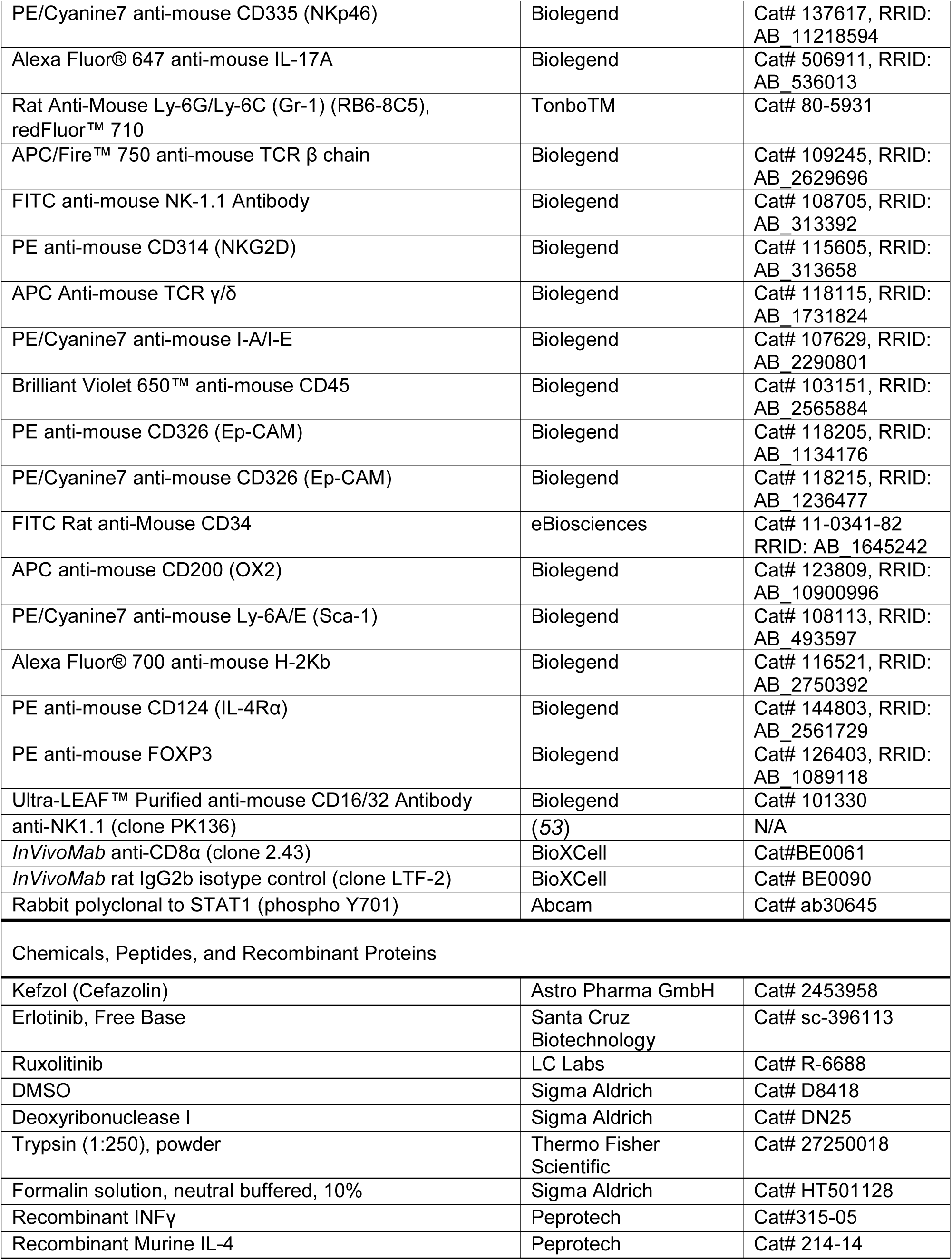
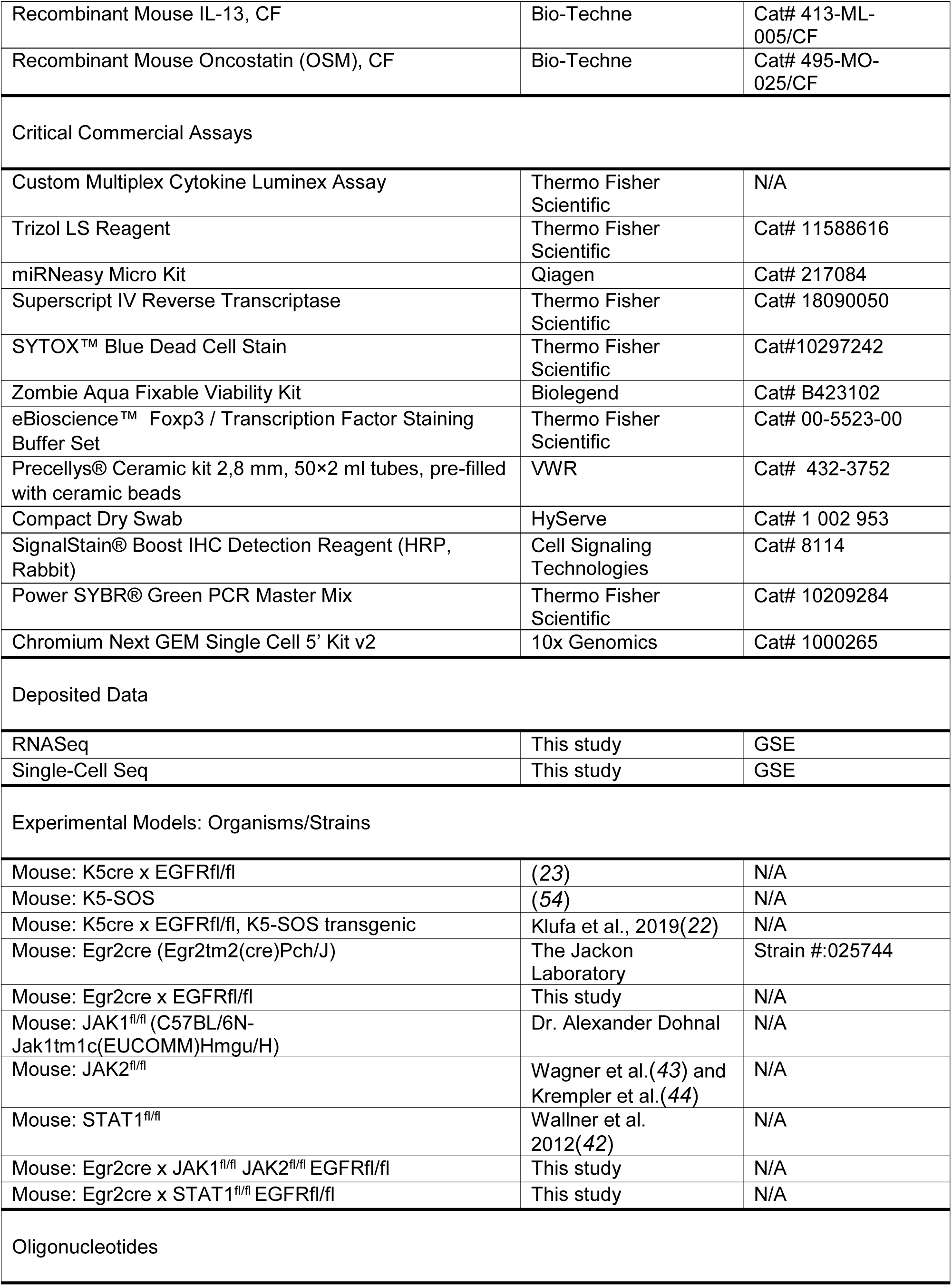
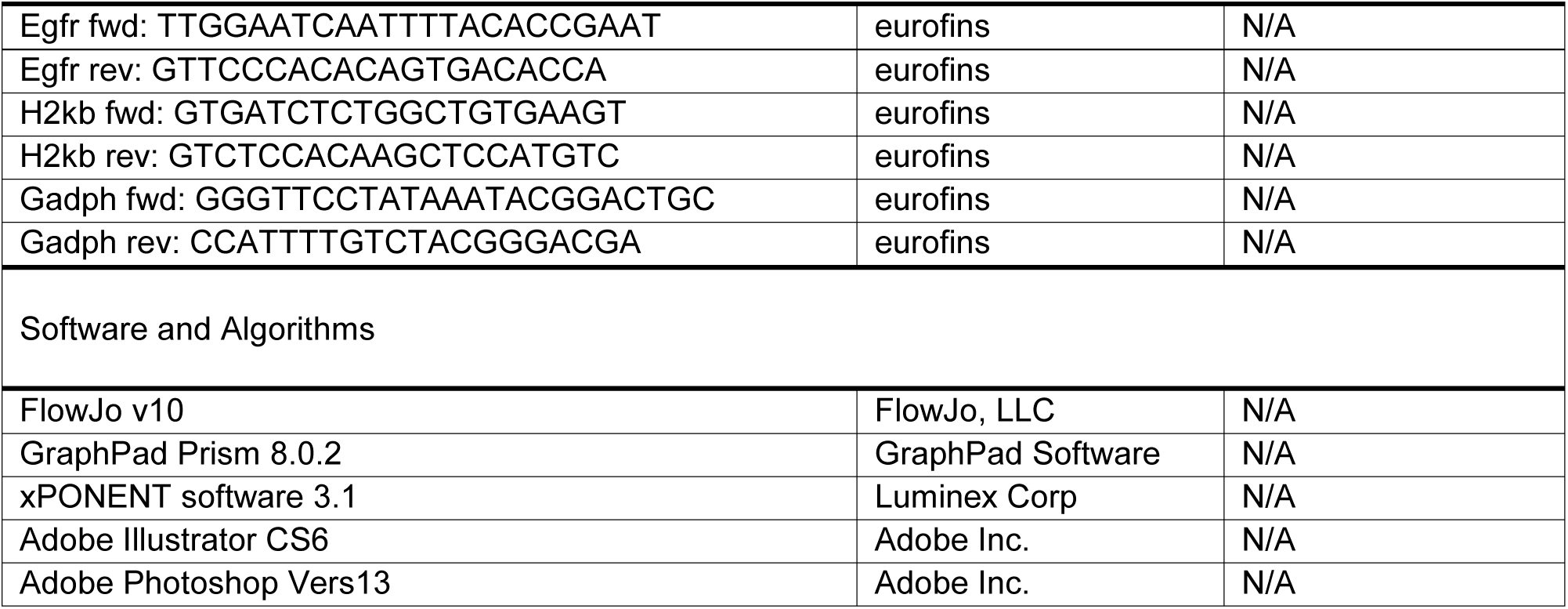

## References

1. M. R. Schneider, R. Schmidt-Ullrich, R. Paus, The hair follicle as a dynamic miniorgan. Curr Biol 19, R132–142 (2009).

2. K. Polak-Witka, L. Rudnicka, U. Blume-Peytavi, A. Vogt, The role of the microbiome in scalp hair follicle biology and disease. Exp Dermatol 29, 286–294 (2020).

3. R. E. Billingham, W. K. Silvers, A Biologist’S Reflections on Dermatology. Journal of Investigative Dermatology 57, 227–240 (1971).

4. R. Paus, N. Ito, M. Takigawa, T. Ito, The hair follicle and immune privilege. J Investig Dermatol Symp Proc 8, 188–194 (2003).

5. J. Agudo et al., Quiescent Tissue Stem Cells Evade Immune Surveillance. Immunity 48, 271–285.e275 (2018).

6. E. C. E. Wang, Z. Dai, A. W. Ferrante, C. G. Drake, A. M. Christiano, A Subset of TREM2^+^ Dermal Macrophages Secretes Oncostatin M to Maintain Hair Follicle Stem Cell Quiescence and Inhibit Hair Growth. Cell Stem Cell 24, 654–669.e656 (2019).

7. Z. Liu et al., Glucocorticoid signaling and regulatory T cells cooperate to maintain the hair-follicle stem-cell niche. Nature Immunology 23, 1086–1097 (2022).

8. W.-B. Wang et al., Developmentally programmed early-age skin localization of iNKT cells supports local tissue development and homeostasis. Nature Immunology 24, 225–238 (2023).

9. N. Ali et al., Regulatory T Cells in Skin Facilitate Epithelial Stem Cell Differentiation. Cell 169, 1119–1129.e1111 (2017).

10. A. Gilhar, Y. Ullmann, T. Berkutzki, B. Assy, R. S. Kalish, Autoimmune hair loss (alopecia areata) transferred by T lymphocytes to human scalp explants on SCID mice. J Clin Invest 101, 62–67 (1998).

11. L. Xing et al., Alopecia areata is driven by cytotoxic T lymphocytes and is reversed by JAK inhibition. Nature medicine 20, 1043–1049 (2014).

12. R. Paus, S. Bulfone-Paus, M. Bertolini, Hair Follicle Immune Privilege Revisited: The Key to Alopecia Areata Management. Journal of Investigative Dermatology Symposium Proceedings 19, S12–S17 (2018).

13. M. J. Harries et al., Lichen Planopilaris and Frontal Fibrosing Alopecia as Model Epithelial Stem Cell Diseases. Trends in Molecular Medicine 24, 435–448 (2018).

14. M. Harries, J. Hardman, I. Chaudhry, E. Poblet, R. Paus, Profiling the human hair follicle immune system in lichen planopilaris and frontal fibrosing alopecia: can macrophage polarization differentiate these two conditions microscopically? British Journal of Dermatology 183, 537–547 (2020).

15. M. J. Harries et al., Lichen planopilaris is characterized by immune privilege collapse of the hair follicle’s epithelial stem cell niche. The Journal of Pathology 231, 236–247 (2013).

16. B. H. Yang et al., A case of cicatricial alopecia associated with erlotinib. Annals of dermatology 23, S350–S353 (2011).

17. B. R. Earl et al., Epidermal growth factor receptor deficiency: Expanding the phenotype beyond infancy. The Journal of Dermatology n/a, (2020).

18. C. W. Franzke et al., Epidermal ADAM17 maintains the skin barrier by regulating EGFR ligand-dependent terminal keratinocyte differentiation. J Exp Med 209, 1105–1119 (2012).

19. J. Nowaczyk, K. Fret, G. Kaminska-Winciorek, L. Rudnicka, J. Czuwara, EGFR inhibitor-induced folliculitis decalvans: a case series and management guidelines. Anticancer Drugs, (2023).

20. M. Holcmann, M. Sibilia, Mechanisms underlying skin disorders induced by EGFR inhibitors. Mol Cell Oncol 2, e1004969 (2015).

21. M. E. Lacouture, Mechanisms of cutaneous toxicities to EGFR inhibitors. Nat Rev Cancer 6, 803–812 (2006).

22. J. Klufa et al., Hair eruption initiates and commensal skin microbiota aggravate adverse events of anti-EGFR therapy. Science Translational Medicine 11, eaax2693 (2019).

23. B. M. Lichtenberger et al., Epidermal EGFR controls cutaneous host defense and prevents inflammation. Sci Transl Med 5, 199ra111 (2013).

24. N. Amberg et al., EGFR Controls Hair Shaft Differentiation in a p53-Independent Manner. iScience 15, 243–256 (2019).

25. F. Mascia et al., Genetic ablation of epidermal EGFR reveals the dynamic origin of adverse effects of anti-EGFR therapy. Sci Transl Med 5, 199ra110 (2013).

26. P. Young et al., E-cadherin controls adherens junctions in the epidermis and the renewal of hair follicles. Embo j 22, 5723–5733 (2003).

27. F. Mascia et al., EGFR regulates the expression of keratinocyte-derived granulocyte/macrophage colony-stimulating factor in vitro and in vivo. J Invest Dermatol 130, 682–693 (2010).

28. A. Limat, T. Wyss-Coray, T. Hunziker, L. R. Braathen, Comparative analysis of surface antigens in cultured human outer root sheath cells and epidermal keratinocytes: persistence of low expression of class I MHC antigens in outer root sheath cells in vitro. Br J Dermatol 131, 184–190 (1994).

29. S. Shao et al., IFN-γ enhances cell-mediated cytotoxicity against keratinocytes via JAK2/STAT1 in lichen planus. Sci Transl Med 11, (2019).

30. B. Homey et al., CCL27-CCR10 interactions regulate T cell-mediated skin inflammation. Nat Med 8, 157–165 (2002).

31. Y. Z. Chiang, C. Bundy, C. E. Griffiths, R. Paus, M. J. Harries, The role of beliefs: lessons from a pilot study on illness perception, psychological distress and quality of life in patients with primary cicatricial alopecia. Br J Dermatol 172, 130–137 (2015).

32. M. Kennedy Crispin, et al., Safety and efficacy of the JAK inhibitor tofacitinib citrate in patients with alopecia areata. JCI Insight 1, e89776 (2016).

33. Y. Tanaka, Y. Luo, J. J. O’Shea, S. Nakayamada, Janus kinase-targeting therapies in rheumatology: a mechanisms-based approach. Nat Rev Rheumatol 18, 133–145 (2022).

34. R. Jerjen, N. Meah, L. Trindade de Carvalho, D. Wall, R. Sinclair, Folliculitis decalvans responsive to tofacitinib: A case series. Dermatol Ther 33, e13968 (2020).

35. A. Moussa, L. Asfour, S. Eisman, B. Bhoyrul, R. Sinclair, Successful treatment of folliculitis decalvans with baricitinib: A case series. Australas J Dermatol 63, 279–281 (2022).

36. C. C. Yang, T. Khanna, B. Sallee, A. M. Christiano, L. A. Bordone, Tofacitinib for the treatment of lichen planopilaris: A case series. Dermatologic Therapy 31, (2018).

37. E. Guttman-Yassky et al., Phase 2a randomized clinical trial of dupilumab (anti-IL-4Rα) for alopecia areata patients. Allergy 77, 897–906 (2022).

38. E. H. C. Wang, et al., Primary cicatricial alopecias are characterized by dysregulation of shared gene expression pathways. PNAS Nexus 1, (2022).

39. C. Tziotzios et al., Genome-wide association study in frontal fibrosing alopecia identifies four susceptibility loci including HLA-B*07:02. Nature Communications 10, 1150 (2019).

40. F. Concha-Benavente, R. M. Srivastava, S. Ferrone, R. L. Ferris, EGFR-mediated tumor immunoescape: The imbalance between phosphorylated STAT1 and phosphorylated STAT3. Oncoimmunology 2, e27215 (2013).

41. R. Pérez-Soler, Can rash associated with HER1/EGFR inhibition be used as a marker of treatment outcome? Oncology (Williston Park) 17, 23–28 (2003).

42. B. Wallner et al., Generation of mice with a conditional Stat1 null allele. Transgenic Research 21, 217–224 (2012).

43. K. U. Wagner et al., Impaired alveologenesis and maintenance of secretory mammary epithelial cells in Jak2 conditional knockout mice. Mol Cell Biol 24(12), 5510–20 (2004).

44. A. Krempler et al., Generation of a conditional knockout allele for the Janus kinase 2 (Jak2) gene in mice. Genesis 40(1), 52–7 (2004).

45. N. Percie du Sert et al., The ARRIVE guidelines 2.0: Updated guidelines for reporting animal research. PLoS Biol 18, e3000410 (2020).

46. N. L. Bray, H. Pimentel, P. Melsted, L. Pachter, Near-optimal probabilistic RNA-seq quantification. Nature Biotechnology 34, 525–527 (2016).

47. M. I. Love, W. Huber, S. Anders, Moderated estimation of fold change and dispersion for RNA-seq data with DESeq2. Genome Biology 15, 550 (2014).

48. Y. Hao et al., Integrated analysis of multimodal single-cell data. Cell 184, 3573–3587.e3529 (2021).

49. M. D. Young, S. Behjati, SoupX removes ambient RNA contamination from droplet-based single-cell RNA sequencing data. GigaScience 9, (2020).

50. P. a. L. Germain, A and Garcia Meixide, C and Macnair, W and Robinson, MD, Doublet identification in single-cell sequencing data using scDblFinder [version 2; peer review: 2 approved]. F1000Research 10, (2022).

51. S. Joost et al., Single-Cell Transcriptomics Reveals that Differentiation and Spatial Signatures Shape Epidermal and Hair Follicle Heterogeneity. Cell Systems 3, 221–237.e229 (2016).

52. S. Jin et al., Inference and analysis of cell-cell communication using CellChat. Nature Communications 12, 1088 (2021).

53. B. Drobits et al., Imiquimod clears tumors in mice independent of adaptive immunity by converting pDCs into tumor-killing effector cells. J Clin Invest 122, 575–585 (2012).

54. M. Sibilia et al., The EGF receptor provides an essential survival signal for SOS-dependent skin tumor development. Cell 102, 211–220 (2000).

