## Supplementary materials for "Scarring hair follicle destruction is driven by the collapse of EGFR-protected JAK-STAT1-sensitive stem cell immune privilege"

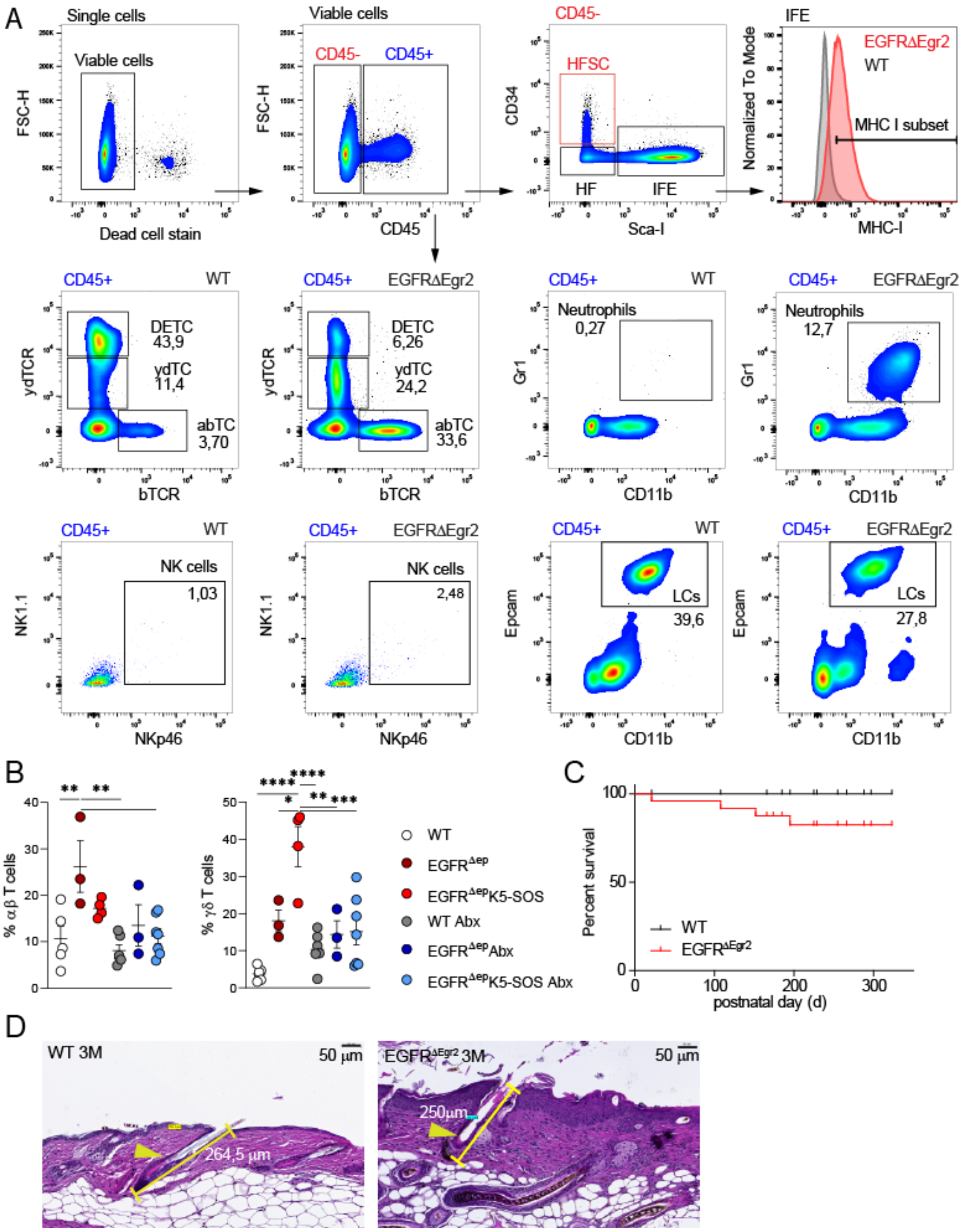


**Figure S1. Hair follicle-specific EGFR deletion induces epidermal immune infiltrate and microbiota-driven inflammatory hair follicle destruction. (A)** Representative pictures of the gating strategy of FACS analysis of epidermal single cell suspensions (WT and EGFR$^{\Delta Egr2}$ mice). CD34$^{+}$ Sca-I$^{-}$ HFSC, CD34$^{-}$ Sca-I$^{-}$ HF and Sca-I$^{+}$ IFE among CD45$^{-}$ keratinocytes. $\gamma\delta$TCR$^{hi}$ dendritic epidermal T cells (DETC), $\gamma\delta$TCR$^{int}$ $\gamma\delta$T cells ($\gamma\delta$TC), $\alpha\beta$T cells ($\alpha\beta$TC), CD11b$^{+}$Gr-1$^{+}$

neutrophils, NK1.1$^+$ Nkp46$^+$ natural killer (NK) cells, CD11b$^+$Epcam$^+$ Langerhans cells (LC) among CD45$^+$ immune cells. **(B)** FACS analysis of $\alpha\beta$T cells and $\gamma\delta$T cells among CD45$^+$ immune cells at 2M. Each dot represents an independent mouse. Mouse models as indicated in the graph. **(C)** Kaplan-Meier survival plot of EGFR$^{\Delta Egr2}$ mice or WT**. (D)** Representative pictures of hair follicle length measured from hematoxylin and eosin (H&E) stained skin sections marked in yellow of WT or EGFR$^{\Delta Egr2}$. Data is presented in ±SEM, *P < 0.05, **P < 0.01, ***P < 0.001 , ****P < 0.0001 by One-Way ANOVA with Tukey's posthoc correction, n≥3.

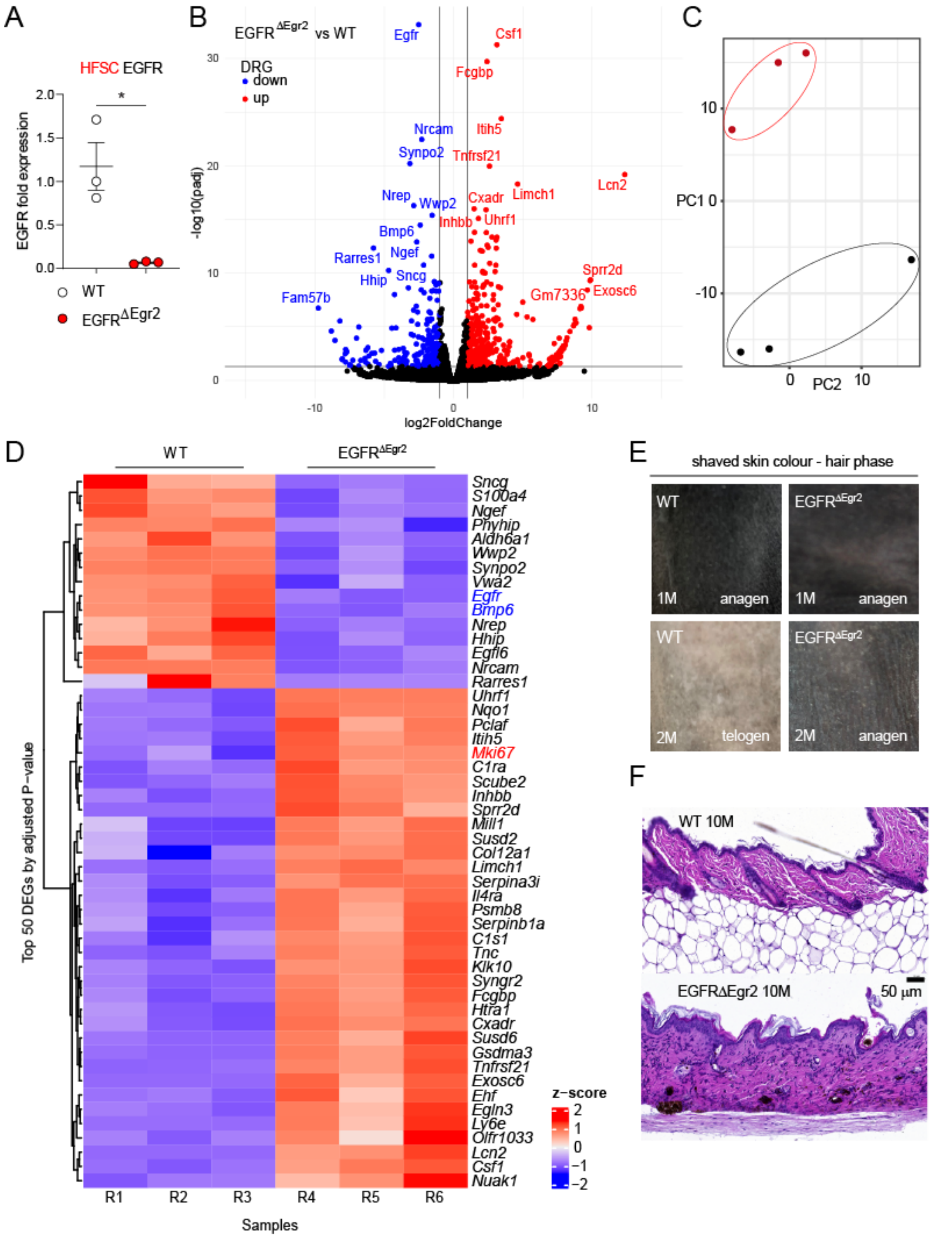


**Figure S2. RNA profiling of FACS sorted EGFR-deficient hair follicle bulge stem cells before scarring hair follicle destruction. (A)** Real-time PCR of EGFR expression of sorted $CD34^{+}ScaI^{-}$ HFSCs of WT and $EGFR^{\Delta Egr2}$ mice at 1M. Data is presented in ±SEM, $^{*}P < 0.05$ by unpaired t-test, n≥3. **(B)** Volcano plot of differentially expressed genes (blue downregulated, red upregulated) of RNA sequencing analysis of $CD34^+$ HFSCs from WT vs $EGFR^{\Delta Egr2}$ mice. Data shown as fold change (log2) and p-value (-log10). **(C)** Principal component (PC) analysis of the RNAseq dataset. **(D)** Heatmap of z-scores of top 50 differentially expressed genes by adjusted p-value of the RNAseq dataset. Genes of special interest are marked in red (up) or blue (down). **(E)** Pictures of shaved backs of WT or $EGFR^{\Delta Egr2}$ at 1M and 2M of age. **(F)** Representative H&E stained skin sections for counting the number of hair follicle units per 1500 μm of WT and $EGFR^{\Delta Egr2}$ skin at 10M of age.

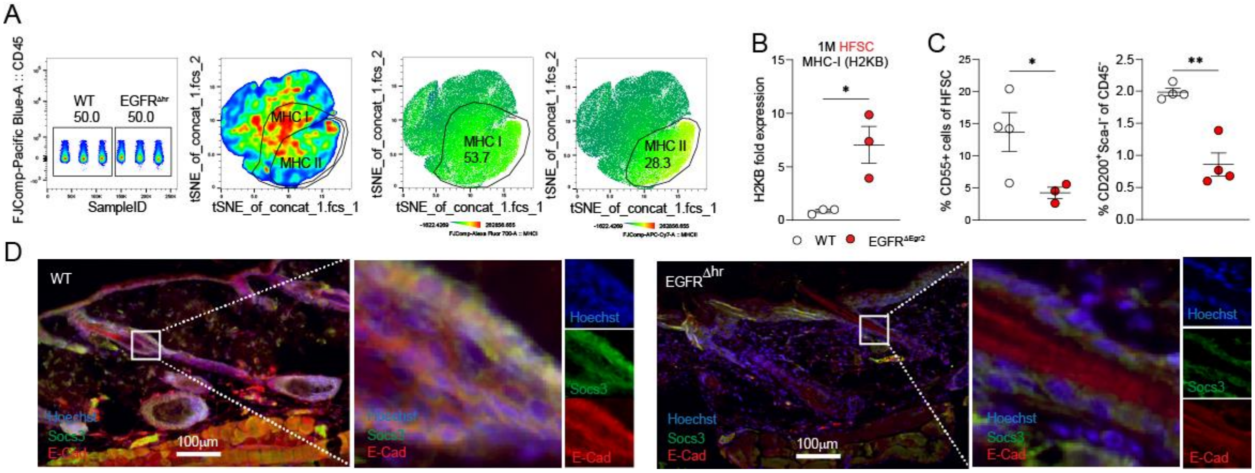


**Figure S3. Immune privilege breakdown indicated by different marker deregulation upon EGFR deletion. (A)** tSNE FACS gating strategy of MHC-I and MHC-II expression among $CD45^-$ keratinocytes of WT or $EGFR^{\Delta Egr2}$ at 5M of age. **(B)** Real-time PCR of MHC-I expression (H2KB) of sorted $CD34^+$ $ScaI^-$ HFSCs of WT or $EGFR^{\Delta Egr2}$ mice at 1M of age. **(C)** CD55 and CD200 surface expression in HFSCs and HF of WT and $EGFR^{\Delta Egr2}$ mice by FACS analysis **(D)** Immunofluorescence staining of E-Cadherin in red and SOCS3 in green of WT and $EGFR^{\Delta Egr2}$ mouse skin sections at 3M. Data is presented in ±SEM, *$P < 0.05$, **$P < 0.01$, ***$P < 0.001$, ****$P < 0.0001$ by unpaired t-test, n≥3.

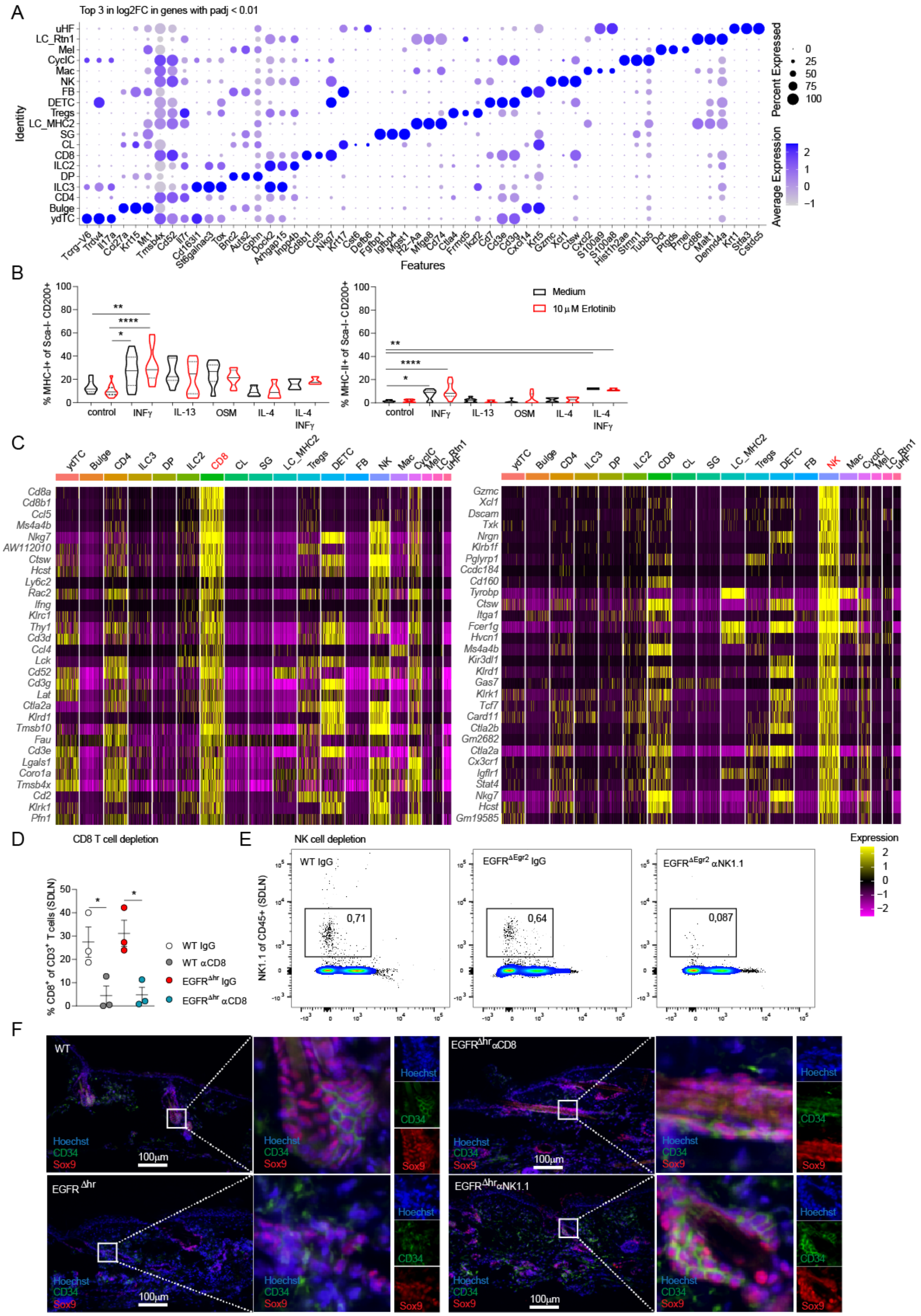

**Figure S4. Single-cell analysis identified hair follicle and immune cell populations in EGFR$^{\Delta Egr2}$ mouse epidermal cell suspensions and cytokine impact of HF specific MHC**

**expression. (A)** Top 3 differentially expressed genes of every cluster of single-cell RNA sequencing analysis of epidermal $CD45^+$ immune cells and $CD45^-$ $Sca\text{-}I^-$ hair follicle cells in $EGFR^{\Delta Egr2}$. **(B)** *Ex vivo* WT skin explants treated with indicated cytokines with or without erlotinib for 48 hours. FACS analysis of MHC-I and -II expressions in $CD200^+$ HF cells. **(C)** Top 30 differentially expressed genes of CD8 and NK cell cluster **(D)** Confirmation of CD8 T cell and **(E)** NK cell depletion by FACS analysis of $EGFR^{\Delta Egr2}$ mice and their respective controls. **(F)** IF staining of $CD34^+$ and $Sox9^+$ stem cells in skin sections of $EGFR^{\Delta Egr2}$ treated with the indicated depletion antibodies and the respective controls. Data is presented in ±SEM, $*P < 0.05$, $**P < 0.01$, $***P < 0.001$, $****P < 0.0001$ by unpaired t-test or One-Way ANOVA with Tukey's posthoc correction, $n \geq 3$.

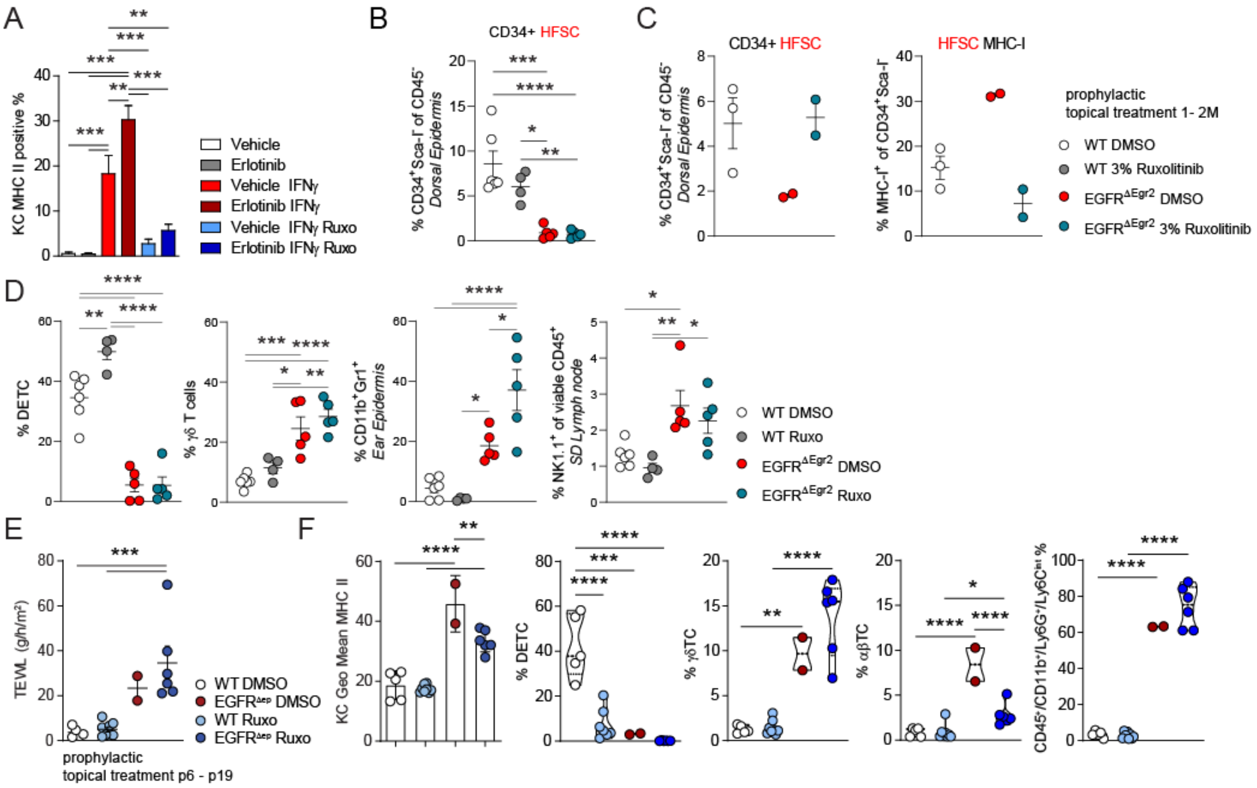


**Figure S5. Therapeutic and prophylactic JAK inhibition ameliorates skin inflammation in $EGFR^{\Delta Egr2}$ and $EGFR^{\Delta ep}$ mice. (A)** MHC-II expression of *in vitro* primary murine KCs of WT mice treated with IFNγ and JAK1/2 inhibitor (ruxolitinib) with or without the EGFR-inhibitor erlotinib. **(B)** Summary of FACS analysis of $CD34^{+}$ hair follicle stem cells from 5M old WT and $EGFR^{\Delta Egr2}$ mice treated therapeutically with DMSO or ruxolitinib in DMSO for 1 month. **(C)** FACS analysis of HFSC and MHC-I expression on $EGFR^{\Delta Egr2}$ and WT mice treated prophylactically with 3% ruxolitinib from 1-2M of age. **(D)** FACS analysis of epidermal $CD45^{+}$ immune cells of $EGFR^{\Delta Egr2}$ and WT treated therapeutically with 3% Ruxolitinib from 5M to 6M. **(e-f)** WT and $EGFR^{\Delta ep}$ mice were treated prophylactically with 3% Ruxolitinib or DMSO starting from P6 until P19 and TEWL was measured from the back-skin (E). FACS analysis of these mice for inflammatory parameters as indicated (F). Data is presented in ±SEM, **$P < 0.01$, ***$P < 0.00$, ****$P < 0.0001$ by one-Way ANOVA with Tukey's posthoc correction, n≥3.

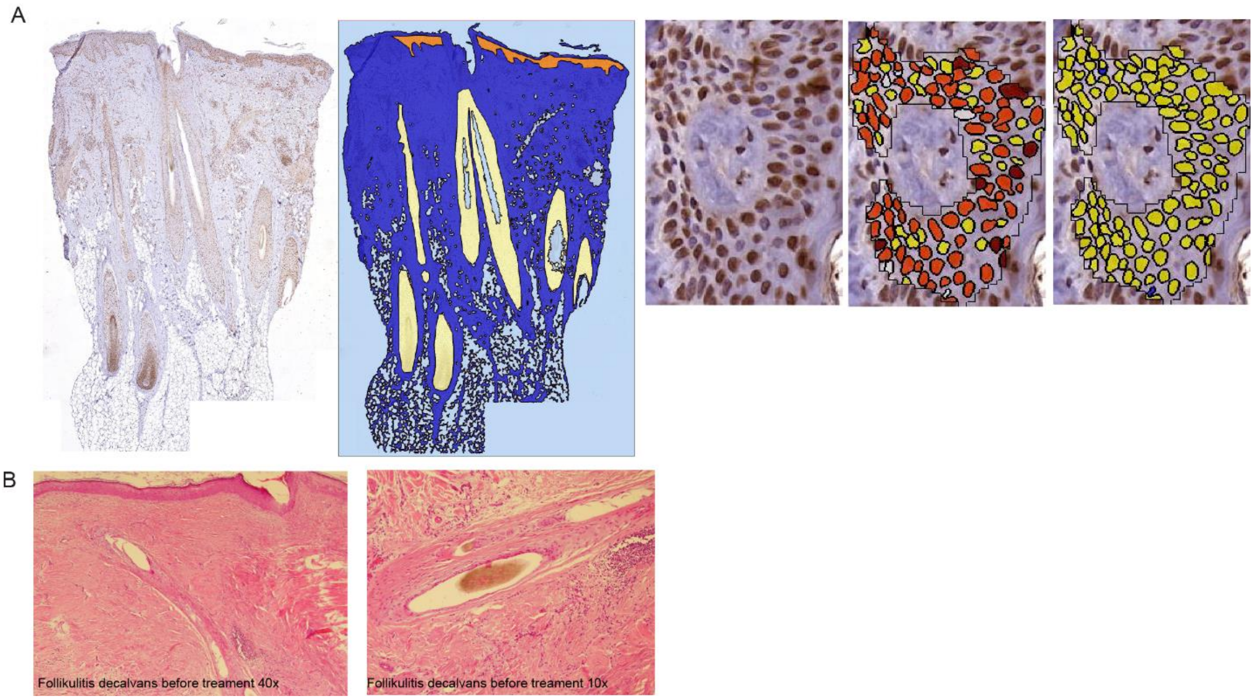


**Figure S6. Quantification of phosphorylated STAT1 in human clinical samples. (A)** Nuclear staining intensity was quantified using definiens software as indicated. Hair follicles (yellow area) and epidermis (orange area) were separately analysed. Medium intensity is shown in red. **(B)** Histopathology of a folliculitis decalvans patient before JAK inhibitor therapy.
